# Ubx orchestrates tissue identity through regional and bidirectional changes to chromatin accessibility

**DOI:** 10.1101/2021.01.15.426863

**Authors:** Ryan Loker, Jordyn E. Sanner, Richard S. Mann

## Abstract

Hox proteins are homeodomain transcription factors that diversify serially homologous segments along the animal body axis, as revealed by the classic bithorax phenotype of *Drosophila melanogaster* where mutations in *Ultrabithorax* (*Ubx*) transform the third thoracic segment into the likeness of the second thoracic segment. To specify segment identity we show that Ubx both increases and decreases chromatin accessibility, coinciding with its role as both an activator and repressor of transcription. Surprisingly, whether Ubx functions as an activator or repressor differs depending on the proximal-distal position in the segment and the availability of Hox cofactors. Ubx-mediated changes to chromatin accessibility positively and negatively impact the binding of Scalloped (Sd), a transcription factor that is required for appendage development in both segments. These findings reveal how a single Hox protein can modify complex gene regulatory networks to transform the identity of an entire tissue.

## Introduction

Among the most famous mutant phenotypes in modern biology is the four-winged ‘bithorax’ fly, in which the third thoracic (T3) segment of *Drosophila melanogaster* is transformed into a nearly complete copy of the second thoracic (T2) segment. This dramatic homeotic transformation of segment identity is caused by loss-of-function mutations in the *Ultrabithorax* (*Ubx*) gene (*1–3*). *Ubx* is one of eight paralogous Hox genes in *Drosophila*, all of which encode homeodomain transcription factors (TFs). Each Hox gene is expressed in a subset of segments along the fly’s anerior-posterior body axis and is responsible for determining their identities. Although the complexity in mammals is compounded by the existence of 39 Hox genes, loss-of-function mutations in the mouse establish that, as in the fly, Hox genes also determine regional identities along the mammalian body axis (*4*).

The dorsal epithelium of wing-bearing T2 and derived T3 segments of *Drosophila* come from the wing and haltere imaginal discs, respectively (Fig. 1A,B). Each disc gives rise to homologous structures of the body wall, hinge, and appendage proper (listed from proximal-most to distal-most position). The dorsal structures of the T3 segment are thought to have evolved from a more ancestral T2-like state (*5*) and are highly modified relative to T2, which largely develops without Hox input (*5–7*). While the wing disc gives rise to the wing and dorsal body wall tissue called the notum, the Ubx-expressing haltere disc gives rise to the haltere, an appendage that beats out of phase with the wings during flight (*8*), and a thin strip of body wall tissue called the postnotum (Fig. 1A,B). Notably, in both imaginal discs the Hox cofactors Homothorax (Hth) and Extradenticle (Exd) are present only in cells that give rise to the body fates and the most proximal parts of the appendages (hinge; see Fig. 1A). Thus, depending on the proximal-distal position, Ubx transforms T2 into T3 both with and without these cofactors (*9, 10*).

**Fig. 1.**
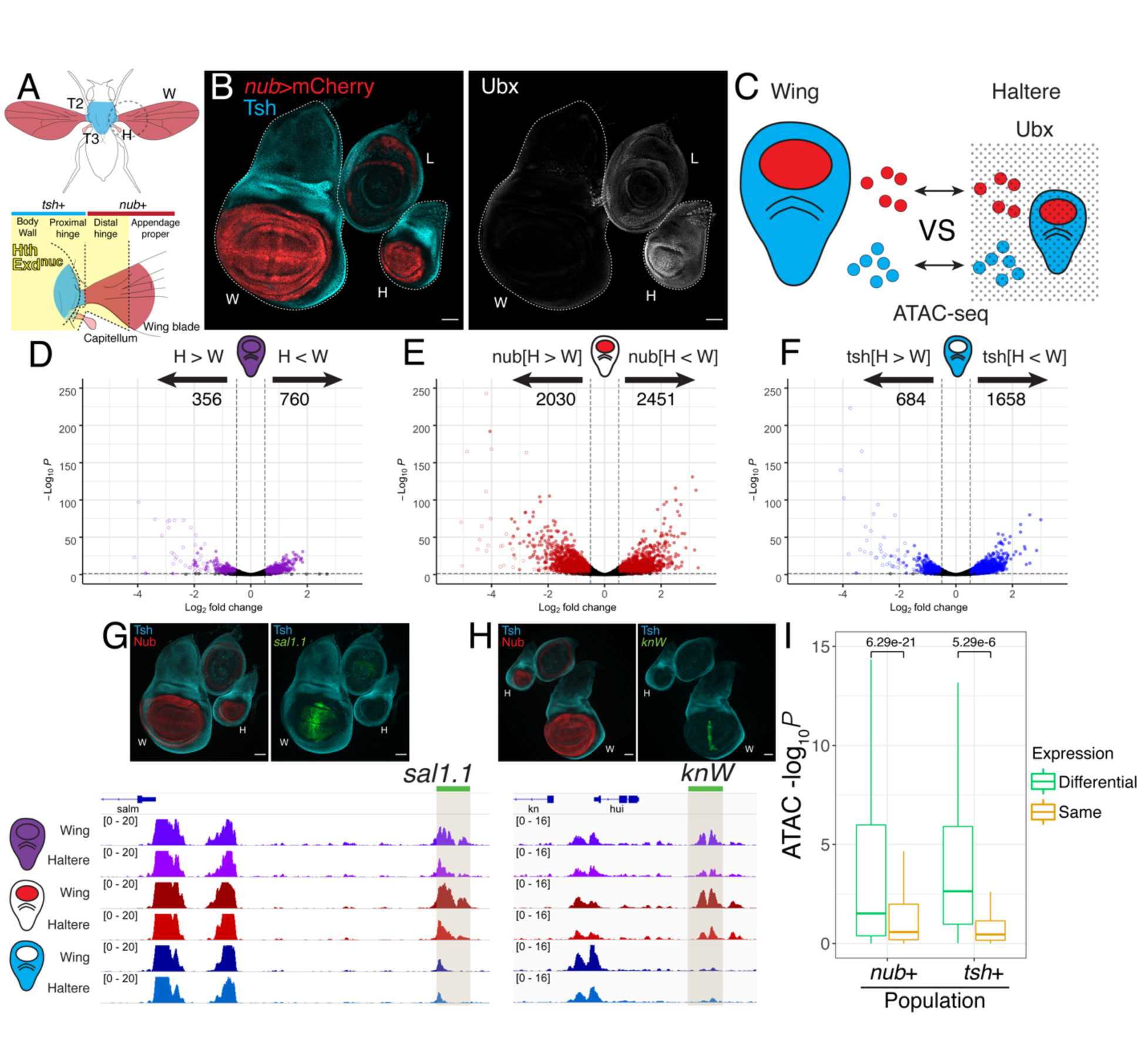
Segment specific chromatin accessibility and gene expression in wing and haltere imaginal discs. A. Schematics of an adult fly highlighting the contributions of the dorsal wing and haltere imaginal discs; the lower panel shows a magnified view of the proximal appendage (hinge) regions. For both the wing-bearing T2 and haltere-bearing T3 segments, blue marks body wall domains (notum and postnotum, respectively) and red marks the appendages (wing and haltere, respectively). The *tsh+* domain (blue) gives rise to the body and proximal hinge, while the *nub+* domain (red) gives rise to the distal hinge and appendage proper (wing blade and capitellum). The Hox cofactor Hth (yellow), which induces the nuclear localization of Exd (Exd^nuc^), is expressed in the body wall, proximal hinge, and distal hinge, but is absent from the appendage proper B. Left, immunostain of 3^rd^ larval instar wing (W) and haltere (H) imaginal discs showing distal (*nub+*, red) and proximal (*tsh+*, blue) populations. Also shown is the T3 leg imaginal disc (L). Right, Ubx is expressed throughout the haltere disc, and is absent from the wing disc. Scale bars for this and subsequent panels are 50uM. C. Experimental scheme to compare chromatin accessibility using ATAC-seq in homologous distal (*nub+*, red) and proximal (*tsh+*, blue) populations of the wing and haltere imaginal discs. Dotted background indicates the presence of Ubx in haltere cells. D-F. Genome-wide comparison of wing and haltere ATAC-seq data for whole tissue (D) *nub+* cells (E) and *tsh+* cells (F). Colored points satisfy a threshold of log2FC>0.5, padj<0.05. Open circles are ATAC peaks within the *Ubx* genomic locus. A common set of 24,915 open chromatin regions, generated by merging ATAC-seq peaks in each sorted data set, was used for comparisons. G-H. ATAC-seq genomic tracks at previously described Ubx target CRMs *sal1.1* (G) and *knW* (H). Cloned fragments driving reporter expression (green) above the genome tracks are indicated by the green bar. I. Comparison of ATAC-seq scores with transcriptome measurements from sorted *nub+* and *tsh+* cells. Differentially expressed genes (DESeq padj<0.01) are significantly more likely to have a differential ATAC peak (DESeq -log_10_pval) compared to genes expressed at similar levels. (See Methods for details.)

Since the original discovery of the *Ubx* mutant phenotype and the realization that Hox genes encode TFs (*11*), many studies have investigated the mechanisms by which Hox proteins activate or repress transcription, largely by analyzing individual *cis-*regulatory modules (CRMs). These studies show that Hox proteins can both activate and repress transcription with and without Hth and Exd (*12*). For example, during embryogenesis Ubx directly represses the leg selector gene *Distaless* (*Dll*) in abdominal segments through a well-characterized CRM that requires Hox-cofactor binding sites (*13, 14*). Ubx also negatively auto-regulates its own expression in the proximal haltere together with Hth-Exd to reduce the levels of its own expression (*10*) and directly represses a handful of targets in the distal-most appendage generating region of the haltere disc without these cofactors (*9, 15–18*). On the other hand, Ubx directly activates CRMs from the *shavenbaby* (*svb*) locus during embryogenesis in conjunction with Hth-Exd (*19*). In light of these and other disparate examples, generalizations for how Hox proteins function *in vivo* have remained elusive. A major barrier has been the technical hurdle of characterizing large sets of Hox-targeted CRMs in specific cell types.

In addition to analyzing individual CRMs, the analysis of chromatin accessibility is a powerful approach to probe the mechanism by which gene regulatory networks are shaped by transcription factors, including Hox proteins (*20–23*). In insect cell culture, embryonic stem cells (ESC), and the mouse limb bud, Hox proteins can increase chromatin accessibility at specific loci in a paralog specific manner (*24–26*). For example the Hox proteins AbdB, HOXC9 and HOX13 paralogs (in *Drosophila* and mammals, respectively) have a relatively high capacity to bind inaccessible chromatin, whereas other Hox factors such as Ubx appear less able to do so and, consequently, are more restricted to binding chromatin that is already in an accessible conformation (*24–27*). Notably, however, Ubx was able to increase accessibility in Kc167 insect cells at a subset of binding sites when it was co-expressed with Hth and Exd, suggesting that the ability to alter chromatin accessibility can be modulated by cofactors (*25*). In a more *in vivo* setting, Hox13 paralogs were shown to increase chromatin accessibility at many sites during the specification of distal fates in the mouse limb bud (*27*). These observations suggest that different potentials to alter chromatin structure may help diversify Hox functions *in vivo*.

In the work described here, we focus on the T2 versus T3 fate decision controlled by Ubx. Instead of comparing different Hox proteins, this system addresses how a single Hox factor modifies the gene regulatory landscape in an entire tissue (T2) to transform it into a different tissue (T3), with homologous cell fates. By carrying out separate chromatin accessibility and transcriptome measurements in the homologous proximal and distal domains of both the haltere and wing imaginal discs, together with profiling Ubx and Hth binding, we show that Ubx both increases and decreases chromatin accessibility at CRMs, coinciding with its role as both an activator and repressor of transcription. Surprisingly, these activities differ depending on the proximal-distal position in the haltere disc. Finally, we demonstrate that Ubx-mediated changes to chromatin accessibility influence the output of another regulatory TF expressed in both T2 and T3 by altering where it binds. Thus, Hox regulation of tissue fate is mediated by regional changes in chromatin accessibility that in turn influence the binding of other TFs, revealing how a single master regulator can transform an entire tissue with multiple cell types.

## Results

### Homologous wing and haltere imaginal discs have distinct chromatin accessibility landscapes

The wing and haltere imaginal discs are comprised of homologous epithelial cell populations and are patterned by a largely conserved network of selector genes and signaling pathways (*18*). We initially performed ATAC-seq to compare the accessible chromatin profiles of the intact wing and haltere imaginal discs at the 3^rd^ larval instar stage. Although the overall profiles are very similar (correlation coefficient = 0.998), consistent with previous observations (*28*), we find 760 sites with decreased accessibility in the haltere (H<W) compared to the wing and 356 sites with increased accessibility (H>W) (DESeq2 (*29*) p-adj<0.05, lfc>0.5; Fig. 1D). Notably, about one fifth of the H>W sites (n=59) are within the *Ubx* locus, and exhibit the highest fold-difference between the wing and haltere discs (Fig. 1D, open circles).

Compared to all accessible regions, differentially accessible sites are more biased to be in introns and intergenic regions (Fig. S1A). All four previously described CRMs regulated by Ubx in the haltere (*15–17*) are identified by this analysis, suggesting that many of the genome-wide differences we identify represent Ubx-mediated changes to the T2 gene regulatory network. The known Ubx-activated and repressed CRMs show an increase and decrease in accessibility in the haltere relative to the wing, respectively, consistent with an inverse relationship between accessibility and repression (*20, 30, 31*) (Fig. S2).

### Chromatin differences downstream of Ubx are regional-specific

Because *Ubx* is expressed in all haltere cells, and must ultimately be responsible for all T3-specific differences, it is possible that the differences in chromatin accessibility measured above exist in all haltere cells, regardless of cell type. Alternatively, Ubx may alter accessibility differently, depending on the cell type. To discriminate between these possibilities, we repeated the ATAC-seq measurements using purified populations of nuclei from homologous distal and proximal domains from the wing and haltere imaginal discs. The distal population, marked by expression of *nubbin* (*nub*), gives rise to the external adult appendages including the distal hinge and appendage proper (wing blade and capitellum for the wing and haltere, respectively) (Fig. 1A,B). The proximal population, marked by expression of *teashirt* (*tsh*), gives rise to the non-appendage thoracic body tissue (notum and postnotum, respectively) and proximal hinge that connects the appendage to the body (Fig. 1A-C).

Comparison of the *nub*+ domains yields 2,451 regions that are less accessible in the haltere compared to the wing (nub[H<W]) and 2,030 regions with increased accessibility in the haltere (nub[H>W]). In the *tsh*+ domain, 1,658 regions have decreased accessibility in the haltere compared to the wing (tsh[H<W]) and 684 have increased accessibility (tsh[H>W]) (Fig. 1D-F). Most of the differential regions identified in the whole disc comparison were also identified in the population-specific comparisons (Fig. S1B). Compared to the whole disc comparisons, the larger number of differentially accessible regions identified in the *tsh+* and *nub+* domains is likely due to greater sensitivity when comparing more homogeneous cell populations.

These data support the idea that changes in chromatin accessibility induced by Ubx are context-specific. Examination of specific CRMs that are differentially expressed in only the *tsh* or *nub* populations further supports this idea. For example, the *sal1.1* and *knW* CRMs (*9, 15, 16*) are repressed by Ubx in *nub+* haltere cells and both enhancers have less accessibility in the *nub+* domain, but no difference in the *tsh*+ domain (Fig. 1G,H).

To assess whether differences in chromatin accessibility correlate with transcriptional changes on a genome-wide scale we performed RNA-seq on the *nub+* and *tsh+* cell populations for both the haltere and wing imaginal discs. Differential analysis performed for *nub+* cells yielded 828 genes downregulated in the haltere and 846 genes with increased expression relative to *nub+* wing cells (Fig. S3A). In the *tsh+* population 126 genes had lower levels and 56 had higher levels in the haltere compared to *tsh+* wing cells (Fig. S3B). For both populations, differentially expressed genes are more likely to have differentially accessible ATAC-seq peaks, compared to genes that are expressed at similar levels (Chi-square test: *P* = 5.99e-47 (nub+) and *P* = 3.06e-36 (tsh+) and Fig. 1I). These results suggest that tissue-specific differences in chromatin accessibility contribute to tissue-specific gene expression.

### Most differences in accessibility are Ubx-dependent

To confirm that the haltere-specific differences in chromatin accessibility depend on Ubx we performed a time-sensitive knockdown of Ubx for 48 hours in the *nub+* domains and repeated the ATAC-seq comparison from the *nub+* population from both the wing and haltere imaginal discs. Following knockdown the Ubx target *salm* is derepressed in the haltere as previously described (*15*) (Fig. 2A) and there is an increase in chromatin accessibility at the *sal1.1* CRM (Fig. 2B), demonstrating that the haltere-specific chromatin accessibility at this locus is dependent on Ubx. Examining the data genome-wide, the majority of tissue-specific differences are lost after knockdown of Ubx. Compared to the wing, 660 regions had less accessibility in the haltere (down from 2,451 in WT) and 237 had increased accessibility (down from 2,030 in WT) (Fig. 2C, S4). Notably, 52 sites in the *Ubx* locus remain more accessible in the haltere in the knockdown. This is expected because, with the exception of autoregulatory elements, the regulation of *Ubx* expression is upstream of Ubx activity and therefore the accessibility of CRMs within *Ubx* should not change in response to reduced Ubx activity. The remaining tissue-specific differences may be due to incomplete knockdown, which is supported by the persistence of weak anti-Ubx antibody staining after knockdown (Fig. 2A).

**Fig. 2.**
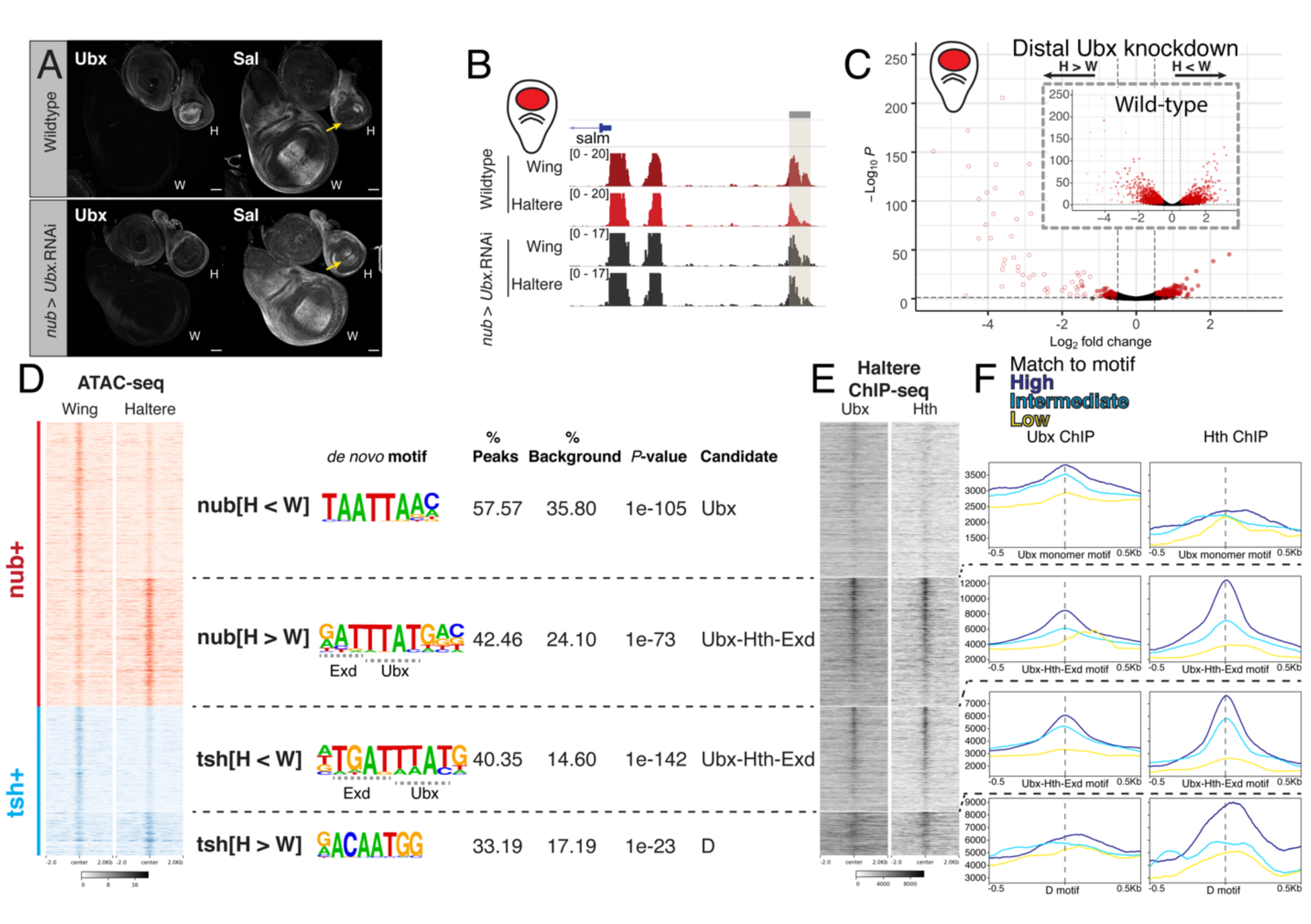
Ubx regulates chromatin accessibility. A. Expression of the Ubx target Salm in wildtype and following Ubx knockdown. De-repression of Salm in the haltere pouch is observed (arrow). B. Genomic tracks showing the *salm* locus. The *sal*1.1 CRM is marked by the grey box. C. Volcano plot comparing *nub+* chromatin accessibility in wing and haltere imaginal discs following knockdown of *Ubx*. Inset; the same comparison in wild type discs is repeated from Fig. 1E for comparison. D. *de novo* motif analysis of the four differential ATAC-seq categories defined in Fig. 1D-F. The top ranked motif for each category is shown. Candidate Ubx and Ubx-Hth-Exd motifs resemble motifs derived from SELEX-seq assays (See Fig. S6A). Heatmaps on the left show the wing and haltere ATAC-seq signals for each of the four categories. E. Heatmap showing the haltere ChIP signal for Ubx and Hth at loci within the differential ATAC-seq categories. Regions are centered around the closest match to the top-ranked *de novo* motif for that category (panel D) and sorted by highest-to-lowest scoring match to that motif. F. Plots showing distribution of average ChIP-signal centered around the same motif as panel E. Each category is split into thirds based on the degree of similarity of motif matching to the top ranked *de-novo* motif for that category. See Methods for details.

### Ubx increases and decreases chromatin accessibility depending on the region of the haltere disc

*De novo* searches for DNA sequence motifs can provide evidence for whether Ubx is directly responsible for changes in chromatin accessibility and if Ubx is binding with or without its cofactors Hth and Exd. Importantly, the *nub+* population includes cells that have these cofactors (*nub+ hth+* cells fated to become the distal hinge) and those that do not (*nub+ hth-* pouch region fated to become the haltere capitellum), while all cells in the proximal *tsh+* population express these cofactors (*tsh+ hth+*; Fig. 1A). Consequently, the association of specific DNA motifs with the gain or loss of accessibility also has the potential to provide spatial information about where Ubx is activating and repressing transcription.

We used an unbiased approach to look for motifs that are enriched in each differentially accessible peak set (nub[H<W], nub[H>W], tsh[H<W], tsh[H>W] (Fig. 2D and S5). Three of the four peak sets contain DNA binding motifs that are predicted to bind Hox proteins as the most enriched sequence. Interestingly, the type of motif differs between peak sets. The nub[H<W] set is highly enriched for a canonical Ubx monomer binding site, suggesting that, as with the previously described *sal*, *knot*, and *ana* targets (*9, 16, 17*), Ubx generally represses transcription as a monomer in the *nub+ hth-* domain. In contrast, the nub[H>W] set is enriched for a motif predicted to bind Ubx in complex with Hth and Exd (Ubx-Hth-Exd motifs (*32*)), suggesting that Ubx activates transcription with these cofactors in the *nub+ hth+* domain. A Ubx-Hth-Exd motif was also enriched in the tsh[H<W] set, suggesting that Ubx represses transcription with these cofactors in the *tsh+ hth+* domain. Equally notable is that none of the sets are enriched for both types of Ubx motifs (Fig. S4). Furthermore, both of the discovered Ubx-related motifs match Ubx and Ubx-Hth-Exd binding sites derived from *in vitro* SELEX-seq experiments, suggesting that they are *bona fide* Ubx monomer and Ubx-Hth-Exd binding sites, respectively (*33, 34*) (Fig. S6). Neither type of Ubx motif is identified in the tsh[H>W] peak set.

Together, these observations suggest that the sign of CRM regulation by Ubx differs depending on where along the proximal-distal axis Ubx functions and whether Hth and Exd are available as cofactors. A corollary to this conclusion is that in each region of the haltere disc Ubx predominantly acts as either a repressor or an activator of transcription. Below we provide additional evidence to support these conclusions by analyzing the activities of specific CRMs as well as the genome-wide binding of Ubx and Hth.

### Ubx binds to CRMs that change chromatin accessibility in the haltere

To further examine the role of Ubx and its cofactors in regulating chromatin accessibility and CRM activity we performed chromatin immunoprecipitation followed by sequencing (ChIP-seq) in whole haltere imaginal discs using antibodies against Ubx and Hth to directly determine their binding profiles genome-wide. The canonical Ubx-Hth-Exd complex motif is the most significantly enriched motif in Ubx ChIP-seq peaks (Fig. S7). The Ubx monomer motif is not significantly enriched in these whole disc ChIP-seq experiments, despite the fact that several CRMs are known to bind Ubx in the absence of cofactors (*9, 16*). Notably, Ubx ChIP-seq experiments in other contexts also failed to identify a strongly enriched monomer motif, possibly reflecting a lower binding affinity or stability of the Ubx monomer to DNA (*24, 35, 36*). For Hth ChIP-seq experiments, both Ubx-Hth-Exd and Hth-Exd binding site motifs are significantly enriched (Fig. S7).

The two ATAC-seq categories that are enriched for the Ubx-Hth-Exd motif (nub[H>W] and tsh[H<W]) both show strong association with Ubx and Hth binding (Fig. 2E). Moreover, the strength of both the Ubx and Hth ChIP signals correlates with the *de novo* enriched Hox-Hth-Exd motif, supporting a direct interaction with these binding sites *in vivo* (Fig. 2F). Although the nub[H<W] category, which is enriched for the Ubx monomer motif, shows generally low ChIP signal for both Ubx and Hth, the strength of Ubx ChIP signal also correlates with the presence of the *de novo* discovered Ubx momomer motif, and the region of maximum binding signal coincides with the location of the motif (Fig. 2F). In contrast, the Hth ChIP signal does not show a similar correlation, supporting the conclusion that Ubx interacts directly with these regions as a monomer without Hth-Exd. Together, these results suggest that Ubx and its cofactors directly bind to many of the sites that have haltere-specific differences in chromatin accessibility. Further, the data suggest that Ubx and Hth binding are directly responsible for many of the observed segment and region-specific differences in chromatin accessibility. The remaining differentially accessible regions that lack motif or ChIP signatures may be indirectly mediated by transcription factors that are downstream of Ubx.

### Differentially accessible loci are composed of tissue-specific CRMs regulated by Ubx

The above analyses suggest that Ubx binds in *nub*+ *hth-* haltere pouch region as a monomer and is associated with a decrease in chromatin accessibility relative to the homologous cells in the wing disc, while Hox-Hth-Exd binding in *nub+ hth+* distal hinge region is associated with greater accessibility. To ask whether these categories reflect true Ubx repressed or activated targets *in vivo* we cloned 20 putative CRMs into reporter constructs and observed their expression pattern in the wing and haltere discs. We chose loci that bind Ubx and have higher or lower accessibility in the *nub+* domain (nub[H>W] and nub[H<W]) in order to ask whether their activity reflects the direction of change in accessibility (activating or repressing, respectively), and whether Ubx regulates them in the predicted region (distal hinge and pouch, respectively, Fig. 3A). We chose candidate CRMs based solely on ATAC-seq differences and Ubx ChIP-seq signal, without taking Hth binding into consideration.

**Fig. 3.**
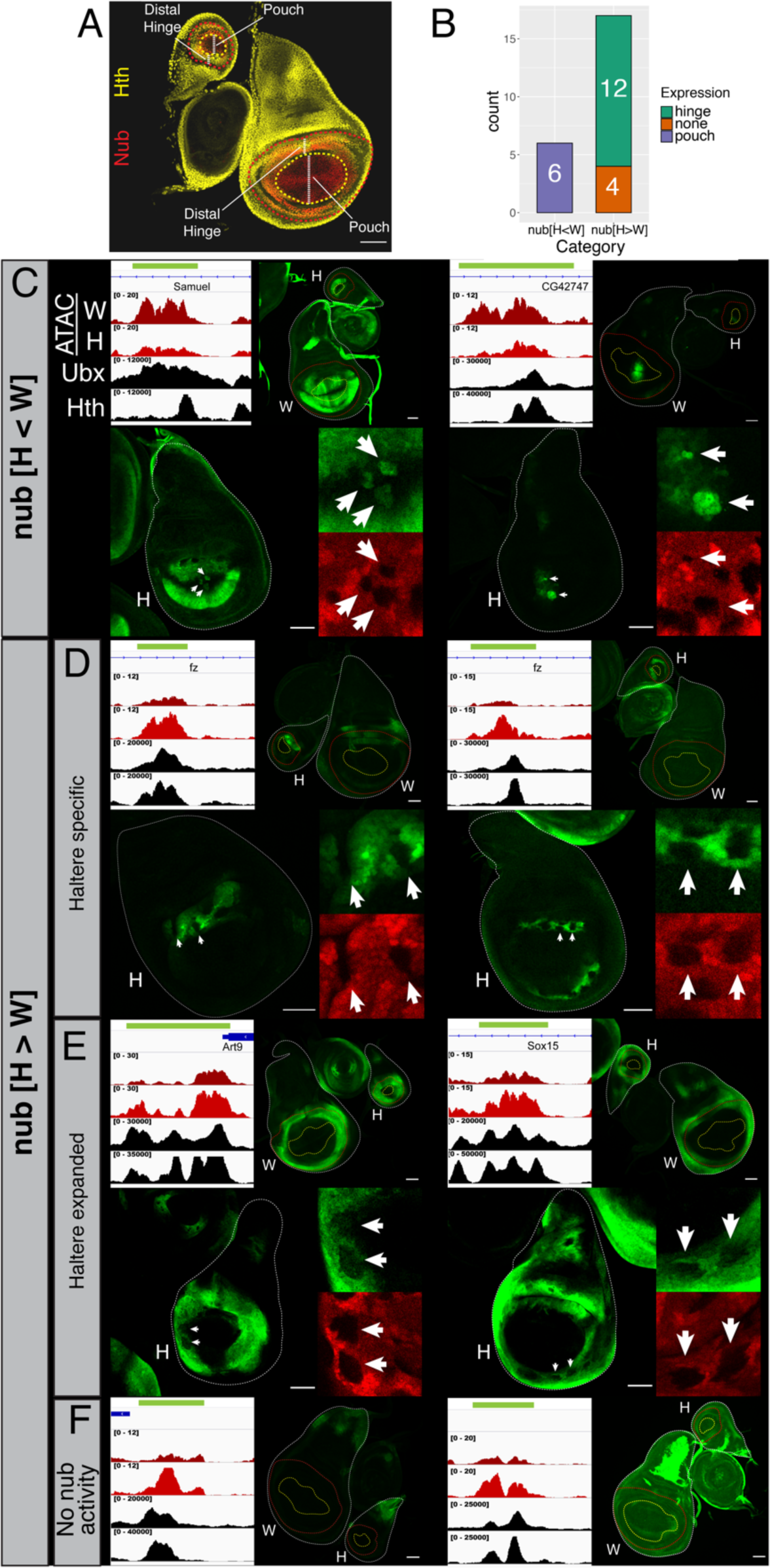
Analysis of Ubx-targeted CRMs. A. Position of homologous distal hinge and pouch domains based on Hth and Nub expression in wing and haltere discs. The proximal edges of Hth and Nub expression are marked with dotted yellow and red lines, respectively. B. Summary of CRM-reporters. The nub[H<W] category includes three previously described CRMs: *sal1.1*, *knW*, and *ana-spot*(*1–3*). C-F. Examples of nub[H<W] and nub[H>W] CRM-reporter genes (green). The upper left panels show genomic tracks for *nub+* ATAC-seq wing, *nub+* ATAC-seq haltere, Ubx ChIP, and Hth ChIP; the upper right panels show wing and haltere disc expression patterns for the reporter genes, and the bottom panels show *Ubx* null somatic clones in the haltere, with a subset of clones magnified in the insets. Clones are marked by the absence of RFP (arrows).

The majority of the cloned regions (3/3 nub[H<W] and 12/17 nub[H>W]) drive reporter expression in a segment-specific manner within the *nub+* domain (Fig. 3B). All three candidates from the nub[H<W] category drive expression in the wing pouch and are less active in the homologous cell population in the haltere (Fig. 3C and S8). These three CRMs behave similarly to the three previously characterized CRMs in the haltere that are repressed by Ubx in this domain, leading to a total of six reporters with similar characteristics (*9, 15–17*) (Fig. 1G-H and S2).

Reporters from the nub[H>W] category (17 in total) are more varied and were grouped into three categories: 1) expressed in the haltere distal hinge domain but not the wing distal hinge (8/17 CRMs, e.g. Fig. 3D and S9), 2) expressed in the distal hinge domains of both tissues, but with a broader pattern in the haltere (5/17 CRMs, e.g. Fig. 3E and S10), and 3) no detectable expression in the *nub+* cells of either disc (4/17 CRMs, Fig. 3F and S11). The third category may represent regions that change accessibility in the 3^rd^ larval instar stage that precede gene expression later in development. Notably, all four of these fragments are active CRMs because they drive expression in other regions of the discs (Figs. 3F and S11). We observed no instances of repression in the distal hinge or activation in the pouch, supporting the conclusion that these Ubx activities are predominantly region-specific.

The differences in reporter activity between the wing and haltere ranged from obvious to subtle. Therefore, for eleven reporters we analyzed mitotic clones of *Ubx* mutant cells in the haltere to confirm that there is a difference in activity downstream of Ubx. In all cases loss of Ubx altered reporter activity toward the wing-like pattern, as expected (Figs. 3C-E, bottom panels and S12). These results strengthen our earlier conclusion that the activity of Ubx as an activator or repressor in the *nub+* cells of the haltere is spatially segregated into the *nub+ hth*+ and *nub+ hth*- domains, respectively. Further, even though they were chosen from the nub[H>W] and nub[H<W] sets, several reporters are also fortuitously expressed in the *tsh+* domain of the wing disc. Further supporting our conclusion that Ubx behaves as a repressor in the *tsh*+ *hth+* domain, in three cases *Ubx*- clones in the haltere derepressed these reporters in that domain (Fig. S13). Added to this list of repressed targets is the autoregulatory *abx* CRM from *Ubx*, which is down-regulated by Ubx-Hth-Exd predominantly in the *tsh+* domain (*10*). Together, these findings reveal that even individual loci can respond to Ubx differently, depending on the region of the imaginal disc, yet they obey the rules uncovered here showing regional differences in Ubx-mediated gene regulation in the haltere disc.

### Changes to chromatin downstream of Ubx alter the binding of another selector TF

How might Ubx-induced changes in chromatin accessibility impact CRM activity and, ultimately, transform T2 into T3? Because both tissues rely on a similar set of patterning TFs, collectively referred to as selector TFs (*37*), we hypothesized that Ubx may either facilitate (in the case of increased accessibility) or prevent (in the case of reduced accessibility) the binding of these shared TFs. As a test of this idea, we focused on the transcription factor Scalloped (Sd) because it has a similar expression pattern in wing and haltere imaginal discs and because it is required for the development of both appendages (Fig. 4A) (*38*). Importantly, Sd is expressed in both the *nub+ hth-* pouch and the *nub+ hth+* hinge domains, where we hypothesize that Ubx is a repressor and an activator, respectively.

**Fig. 4.**
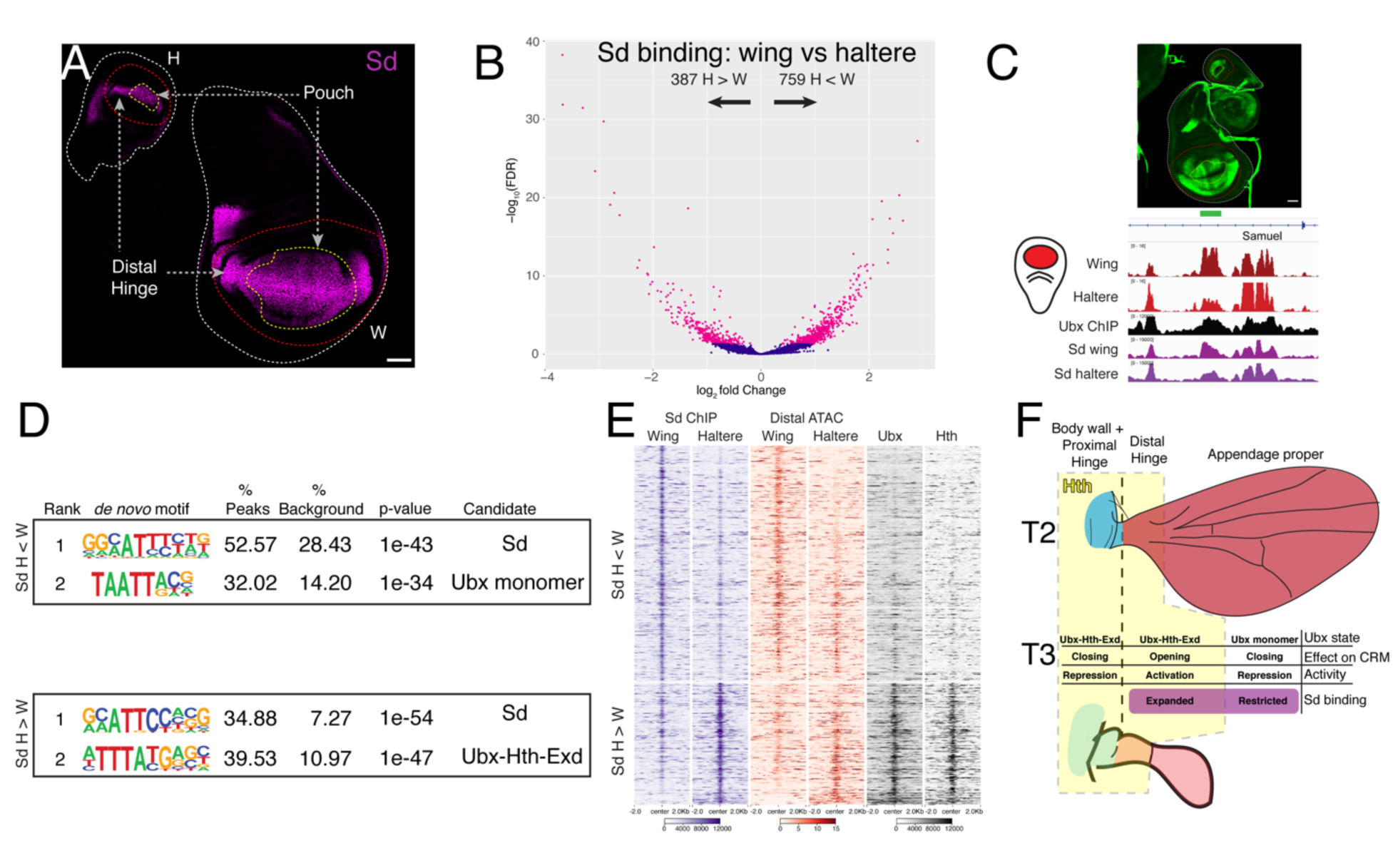
Ubx-mediated changes to chromatin accessibility changes where Sd binds. A. Homologous patterns of Sd expression in the wing and haltere imaginal discs. In both tissues Sd is expressed in the pouch, in the distal hinge and along the dorsal-ventral compartment boundary. Boundaries of Nub (Red) and Hth (Yellow) expression are indicated with dotted lines as in Fig. 3. B. Volcano plot comparing Sd binding in wing and haltere imaginal discs (Diffbind FDR<0.05). C. Genomic tracks near the *Samuel* CRM (green box) and reporter expression driven by this CRM in wing and haltere discs. D. *de novo* motif analysis of the disc-specific Sd binding peaks for the Sd H<W and H>W categories. E. Heatmaps showing the ChIP signal for differential Sd binding, *nub+* ATAC-seq signal, Ubx ChIP signal, and Hth ChIP signal. Regions are sorted based on highest-to-lowest W:H ratio of Distal ATAC-seq signal at the peak center. The top set shows the Sd H<W category and the bottom set shows the Sd H>W category as defined in panel B. F. Summary defining the three domains in T2 and T3, whether Ubx acts as a monomer or Ubx- Hth-Exd complex, whether Ubx opens or closes chromatin, and the effect on Sd binding.

To ask whether Sd binding differs in the haltere and wing imaginal discs, we performed ChIP-seq for Sd in both discs and compared the binding pattern. A subset of Sd binding sites (8.3%) are disc-specific: 387 peaks show stronger binding in the haltere, while 759 peaks are stronger in the wing (Fig. 4B). *De novo* motif searches around both sets of Sd binding peaks show that, in addition to canonical Sd motifs, Ubx motifs are enriched to similar levels, suggesting that they are also targeted by Ubx (Fig. 4D). However, as with the ATAC-seq data, the type of Ubx motif is distinct in peaks biased towards the different discs. H>W Sd binding events are enriched for the Hox-Hth-Exd motif, while H<W Sd binding is associated with Ubx monomer motifs. Additionally, of the 661 peaks that have both tissue specific Sd binding and differences in chromatin accessibility (58% of all tissue-specific Sd binding), 229 (35%) have a Ubx ChIP peak (Fig. 4E and S14, and see Fig. 4C for a specific example at the *Samuel* CRM). These data suggest that the binding of Sd is responsive to the presence of Ubx locally at the CRM, and points to a potential mechanism for how Ubx alters the output of shared TFs both positively and negatively: Ubx binding to monomer sites reduces accessibility and inhibits Sd binding in the pouch, while Ubx binding to Ubx-Hth-Exd sites increases accessibility and facilitates Sd binding in the distal hinge.

## Discussion

We have performed a series of experiments to probe the mechanism by which the Hox protein Ubx transforms the wing-bearing T2 segment into the homologous haltere-bearing T3 segment of the adult fly. We find that in a single tissue with multiple cell types, Ubx has diverse effects on *cis*-regulatory chromatin and ultimately gene regulation, functioning as both an activator and repressor of transcription. Unexpectedly, the data suggest that Ubx functions in three distinct modes that are spatially segregated in the imaginal disc: 1) Ubx reduces chromatin accessibility and represses transcription in the distal-most pouch domain as a monomer, 2) Ubx increases accessibility and activates transcription in complex with Hth-Exd in the distal hinge, and 3) Ubx reduces accessibility and represses transcription in the body wall and proximal hinge in complex with Hth-Exd. Below we discuss these findings in the context of previous research and propose general mechanisms for Hox-mediated gene regulation.

### Ubx both increases and decreases chromatin accessibility

Our data are consistent with the idea that when bound as a monomer, Ubx predominantly functions as a transcriptional repressor associated with decreasing chromatin accessibility, but when bound as a Ubx-Hth-Exd complex, Ubx can both activate and repress gene expression depending on the cell context. The finding that the Ubx-Hth-Exd complex has different properties from the Ubx monomer is consistent with experiments carried out in Kc167 cells, where Ubx was also able to increase chromatin accessibility only when co-expressed with its cofactors (*25*). However, in addition to increasing accessibility, we show that Ubx-Hth-Exd also has the potential to decrease accessibility in a different context, in cells that give rise to the body and proximal hinge of the T3 segment. Additionally, in contrast to our findings, no Hox protein was previously shown to decrease chromatin accessibility in the Kc167 or other experimental systems. Our ability to observe both increases and decreases in chromatin accessibility may be a consequence of studying the T2 versus T3 binary transformation, which includes multiple cell fates that are all modified by the presence of the same Hox protein. This is in contrast to previously characterized Hox-regulated systems, such as the vertebrate limb bud (*27*) or the induction of motor neuron fates from ESCs (*26*), where instead of transforming one tissue into another, Hox proteins promote the development of specific cell fates from less differentiated progenitors.

The precise nature of how Ubx alters chromatin accessibility requires further investigation. One potential mechanism involves the recruitment of chromatin modifying factors, several of which have been shown to interact with Hox proteins (*39, 40*), to modulate the compaction of the local CRM structure as suggested for the repression of *Dll* by Ubx (*41*). Alternatively, Ubx may compete with the binding of activator TFs (in the case of haltere repression) or facilitate activator binding through nucleosome-mediated cooperativity (*42*) (in the case of haltere activation). Notably, in the cases of Ubx repression of *knot* (*16*) and *Dll* (*13, 14*), the repressive binding input of Ubx into the relevant CRMs was shown to be separable from the activating input from other TFs, suggesting, at least for these cases (which involve monomer and Ubx-Hth-Exd input, respectively) competition for binding is unlikely to be involved.

### Spatial determinants of Hox activity

Although Hox proteins can function as both activators and repressors of gene expression, it is generally unknown if, for a particular TF, whether both activities concurrently operate in the same cell, or whether they perform only one activity depending on the cell type. The first scenario implies that each target CRM recruits a different set of TFs based on the composition of binding sites, and that the combination of the recruited factors, termed collaborators, determines the regulatory outcome (*12, 13, 15, 43*). The failure of *de novo* motif searches to identify motifs enriched to a similar degree as Hox-related ones in our ATAC-seq peak sets suggests that there is not a single or small number of collaborator TFs that work with Ubx within each region of the haltere disc, consistent with observations that Hox proteins can interact with many TFs (*44, 45*). However, because Ubx activity appears to be largely constrained to either activation or repression in each region of the haltere disc, collaborators are unlikely to be the sole determinant of the sign of gene regulation, as it would require all collaborators that function in the same region of the disc to act as either activators or repressors.

The alternative scenario, in which Hox proteins execute the same regulatory output in a given cell type, implies that Hox proteins exist in distinct cell-type specific regulatory complexes that function either as dedicated activators or repressors. The three spatially segregated domains of the haltere disc identified here in which Ubx acts mainly as an activator or repressor, together with the hundreds of similarly regulated CRMs in each of these domains, provide strong support for this scenario. In this case, collaborator TFs may refine in which subset of cells within a haltere domain each CRM is regulated by Ubx, perhaps by recruiting or stabilizing Ubx binding, or tuning the strength of the regulation in a particular direction. Consistent with this notion, the expression patterns of many genes are modified in the haltere relative to the wing in only part of the disc. For example, *wingless* is repressed by Ubx only in the posterior compartment of the pouch domain (*18*). Another example is the *Samuel* CRM identified here, which is repressed by Ubx in a subset of pouch cells (Fig. S15).

### Hox proteins influence the output of other selector TFs by altering where they bind

As serially homologous tissues, the wing and haltere imaginal discs have a very similar organization of spatially restricted signaling pathways and share many of the same regionally expressed TFs. For example, in both discs the secreted morphogen Decapentaplegic (Dpp) is expressed along the anterior-posterior compartment boundary, and the selector TFs Engrailed (En) and Sd are expressed in homologous posterior and appendage-generating compartments, respectively (*18*). Ubx operates upon this common ground-state to modify how these shared pathways and selector TFs are deployed in a T3-specific manner (*18, 46, 47*). Ubx may alter the output from these shared systems by modifying the expression of the signaling molecules themselves, such as *wingless* repression in the posterior compartment of the haltere disc (*18*), or by modifying the distribution of secreted signals, as in the case of Dpp signaling (*46*). Our results reveal that Hox proteins can also modify the output of shared regulators by altering where they bind through changes to *cis*-regulatory chromatin accessibility. In the distal hinge, Ubx, together with Hth-Exd, facilitates Sd binding to sites that are less accessible in the wing, consistent with a pioneer-like activity for the Ubx-Hth-Exd complex in this region of the disc. A pioneering role for the mammalian HOX13 paralogs, HOXA13 and HOXD13, was also proposed during specification of the distal limb bud (*27*). In this example, HOX13 paralogs were required for a subset of open chromatin regions within these cells, and could expand binding of another HOX protein, HOXA11, when it was ectopically expressed. In the haltere Ubx also functions in the opposite direction to restrict access of TFs to CRMs: cofactor-independent Ubx activity decreases accessibility and blocks Sd binding. We suggest that Ubx is likely to both facilitate and block the ability of other shared regulators to bind DNA, revealing a novel mechanism for diversifying homologous cell fates in T3. More generally, we speculate that this mechanism is used by other Hox proteins when they modify serially homologous tissue fates.

### Diversification of body structures by Hox proteins

We have found that Ubx alters chromatin accessibility to modify the ground-state T2 segment into the more derived T3 segment, with its unique body wall and appendage morphologies. An interesting trend emerges from comparison of the homologous adult structures that are modified by Ubx via activation or repression. In the capitellum and notum/proximal hinge, where Ubx-mediated gene repression dominates, the T3 morphology, broadly characterized, has both a reduced size and complexity. The latter can be observed through the loss of characteristic features of the T2 appendage and notum, such as highly patterned veins and large bristles (called macrochaetes), respectively. In contrast, the distal hinge of the haltere, where Ubx-mediated gene activation is the rule, develops complex T3-specific structures that are required for the haltere to provide critical sensory feedback during flight, most notably via arrays of mechanosensory neurons (*8*). We speculate that diversification of tissue morphology by Hox proteins may follow a pattern wherein repressive activities contribute to simplifying the morphology of a tissue while gene activation may be required to generate novel complex cell types.

## Acknowledgements

We thank Roumen Voutev for providing reporter plasmids, Vikki Weake for providing the UAS-Kash.GFP strain, Chaitanya Rastogi and Harmen Bussemaker for helping to deploy SELEX-seq models genome-wide, Siqian Feng for advice on ChIP-seq, Kevin White for Ubx and GFP antibodies, Stavros Lomvardas for sharing equipment, and the Bloomington Drosophila Stock Center for fly stocks. We are also grateful to David Stern, Stavros Lomvardas, and Rebecca Delker for detailed comments on the manuscript. This work was supported by NIH grant R35GM118336 awarded to R.S.M.

## Author Contributions

R.L: Conceptualization, Methodology, Investigation, Formal analysis, Visualization, Writing – Original Draft, Writing – Review&Editing. J.S: Investigation. R.S.M: Conceptualization, Methodology, Writing – Original Draft, Writing – Review&Editing, Funding acquisition

## Data and materials availability

All data generated in this study will be deposited in the GEO database

## Materials and Methods

### *Drosophila* strains and transgenes

Transgenes used in this study are as follows:

*nub*.Gal4 (*48*)

*tsh*.Gal4 (*48*)

UAS.Kash-GFP (Gift of Dr. Vikki Weake, Purdue Univ.) (*49*)

Sd-GFP (protein-trap fusion, FlyTrap, Bloomington # 50827) (*50*)

UAS.Ubx.RNAi (chr. II) (*10*)

UAS.Ubx.RNAi (chr. III) (*10*)

UAS-mCherry.nls (Bloomington # 38425)

tub.Gal80^ts^ (*51*)

### Construction of enhancer reporter genes

Genomic fragments corresponding to putative CRMs were amplified from a generic laboratory *yw* stock. Regions were placed via restriction-mediated cloning into the multiple cloning site of pRVV54 (*52*), in which the *lacZ* ORF was replaced with the eGFP ORF (gift of Roumen Voutev, Columbia University). Coordinates of selected regions are described in Table S1. CRMs of *sal1.1* and *knW* were cloned using coordinates previously described (*16, 53*). *sal1.1* was inserted into pRVV54-LacZ and *knW* was synthesized as a full length fragment by Genewiz and inserted via restriction digest into a pH-Stinger (*54*) plasmid in which a attB sequence was inserted into the AatII restriction site. All reporters were integrated into the genome using PhiC31 system (*55*) at the Attp2 landing site. Primers used for each reporter are listed in Table S1.

### Clonal analysis

*Ubx* mitotic null clones were made using the Flp/FRT system (*56*) using the null *Ubx^9-22^* allele (*57*). Larvae were heatshocked at 37°C for 40-50 minutes at the end of the 2^rd^ instar stage and analyzed 48 hours later.

### Immunohistochemistry

Wandering 3^rd^ instar larval heads were dissected and inverted in PBS, followed by fixation in 4% PFA for 25 minutes at room temperature. Heads were then washed 2X 30 minutes in staining solution (SS: PBS, 1% BSA, 0.3% Triton-X). Primary antibodies were then added for incubation overnight in SS at 4°C. Heads were washed 4X 10 minutes in SS and incubated with fluorescent secondary antibodies for 2 hours at room temperature in dark, and washed as before. Heads were incubated overnight in Vectashield containing DAPI, and imaginal discs were subsequently dissected and mounted for imaging using a confocal microscope (Leica SP5 or Zeiss LSM 800) Primary antibodies used were:

anti-Ubx (Mouse, FP3.38(*57*), Developmental Studies Hybridoma Bank)

anti-Hth (Guinea Pig, Gp115, produced by Calico)

anti-Tsh (Guniea pig, Gp68, produced by Calico)

anti-Nub (Mouse, 2D4(*58*), Developmental Studies Hybridoma Bank)

anti-Spalt (gift from James Hombría, CABD)

anti-GFP (Goat, gift from Kevin White, UChicago)

anti-β-galactosidase (Rabbit, Cappel)

### Nuclei sorting

Nuclei were magnetically sorted from wing and haltere imaginal discs using the UAS.Kash-GFP transgene as previously described (*49*) with slight modifications. Briefly imaginal discs were isolated from larvae at the 3^rd^ instar wandering stage in PBS with 0.01% tween-20 on ice. Dissected tissue was then washed 2X in chilled nuclei extraction buffer (NEB: 10 mM HEPES, pH =7.5; 2.5 mM MgCl; 10 mM KCl). Nuclei were extracted in a 1 mL dounce on ice using 20 strokes of the loose pestle, followed by a 10-minute incubation, and 25 strokes of the tight pestle. Nuclei were then filtered over 30 uM cell filter, and pre-cleared for 10min with 5ul of Protein-G Dynabeads in NEB supplemented with 0.1% tween-20. Pre-clearing beads were removed with a magnet and nuclei were added to a new tube containing anti-GFP coated Dynabeads and incubated with rotation for 30 min at 4°C. Afterwards bead-bound nuclei were washed 4X (5min each) with nuclei wash buffer (15 mM TRIS, pH=7.5; 50 mM NaCl; 40 mM KCl; 2 mM MgCl2; 0.1 % Tween-20). Isolated nuclei were counted on a hemocytometer, and used for ATAC-seq or RNA-seq.

### ATAC-seq library preparation and sequencing

ATAC-seq was performed on 50,000 nuclei as previously described (*59*). Libraries were sequenced using a 150-cycle high output (wild-type samples) or 75-cycle high output (RNAi samples) with paired end sequencing using an Illumina Nextseq.

### RNAi knockdown

Larvae of the genotype *yw; nub.G4, tub-Gal80^ts^/UAS.Ubx.RNAi; UAS.Kash-GFP/UAS.Ubx.RNAi* were raised at 18°C until early 3^rd^ instar and subsequently shifted to 29°C to permit expression of RNAi for 48 hours. Wandering 3^rd^ instar larvae were collected, and subjected to ATAC-seq as described above, separately for wing and haltere discs.

### RNA-seq library preparation and sequencing

RNA was extracted from sorted nuclei using TRIzol and purified using the Zymo Direct-zol RNA Microprep kit. RNA-seq libraries were prepared using total rRNA depleted RNA using Nugen Ovation Drosophila RNA-seq system. Libraries were sequenced using a 150-cycle high output with paired end sequencing using an Illumina Nextseq.

### ChIP-seq library preparation and sequencing

ChIP-seq using wing and haltere imaginal discs was performed as described previously (*60*) with minor modifications according to (*61, 62*). 3^rd^ instar larval heads were dissected and inverted in PBS on ice. Heads were fixed for 20 minutes in 1.8% PFA in crosslinking medium (10 mM HEPES, pH=8.0; 100 mM NaCl; 1 mM EDTA, pH=8.0; 0.5 mM EGTA, pH=8.0) at room-temperature with rotation, and subsequently quenched (Quench solution: 1xPBS; 125 mM glycine; 0.1% Triton X-100). Fixed-heads were then washed 2X in buffer A (10 mM HEPES, pH=8.0; 10 mM EDTA, pH=8.0; 0.5 mM EGTA, pH=8.0, 0.25 % Triton X-100) and 2X buffer B (10 mM HEPES, pH=8.0; 200 mM NaCl; 1 mM EDTA, pH=8.0; 0.5 mM EGTA, pH=8.0; 0.01% Triton X-100) 10 minutes each at 4°C. Wing or haltere discs were then dissected and placed in sonication buffer (10 mM HEPES, pH = 8.0;1 mM EDTA, pH = 8.0; 0.5 mM EGTA, pH = 8.0, 0.1 % SDS). Chromatin sonication was performed using a Covaris S2 instrument at settings (105W; 2 % Duty; 15 minutes).

Sonicated chromatin was brought to 1X mild-RIPA (10 mM Tris-HCl, pH=8.0; 1 mM EDTA, pH=8.0; 150 mM NaCl; 1% Triton X-100) concentration and pre-cleared with Dynabeads for 1 hour at 4°C with rotation. Pre-clearing beads were removed and antibody was added for incubation overnight. Dynabeads were added and incubated for 3 hrs. Bead bound antibody-chromatin complexes were washed as follows 2X RIPA LS (10 mM Tris-HCl, pH=8.0; 1 mM EDTA, pH=8.0; 150 mM NaCl; 1% Triton X-100; 0.1 % SDS; 0.1 % DOC), 2X RIPA HS (10 mM Tris-HCl, pH=8.0; 1 mM EDTA, pH=8.0; 500 mM NaCl; 1% Triton X-100; 0.1 % SDS; 0.1 % DOC), 1X LiCl (10mM Tris-HCl, pH=8.0; 1 mM EDTA, pH=8.0; 250 mM LiCl; 0.5 % IGEPAL CA-630; 0.5 % DOC), 1X TE (10 mM Tris-HCl, pH=8.0; 1 mM EDTA, pH=8.0). Samples were then treated with RNAse and proteinase K, and chromatin was isolated using phenol-chloroform.

Antibodies used were anti-Ubx (7701 (*63*), 1:100 dilution for IP, gift from Kevin White, U. Chicago), anti-Hth (Gp52 (*64*), 1:300 dilution for IP), and anti-GFP (used for Sd-GFP; ab290, Abcam; 1:300 dilution for IP).

ChIP-seq libraries were made following the NEBnext UltraII kit (NEB) and associated protocol. Libraries were sequenced using a 75-cycle high output with single end sequencing using an Illumina Nextseq.

### ATAC-seq data processing

Reads were mapped using Bowtie2 to the dm6 genome assembly. Mapped reads were then filtered for map quality (SAMtools (*65*)) and duplicates (Picard MarkDuplicates). The Galaxy platform (*66*) was used for these pre-processing steps.

Genome-track files were created using Deeptools (BamCoverage; RPGC normalization).

Differential analysis was performed using DESeq2 (*29*) on a common interval of 24,915 peaks generated by merging ATAC-seq peaks called by MACS2 (*67*)(--nomodel --call-summits) from the all wild-type sorted data sets. Cut off used for calling differential accessibility was Log_2_Fold change > 0.5 and adjusted p-value (padj) < 0.05. Peaks within the extended *Ubx* genomic locus were defined by (Chr3R: 16655898- 16807343). Heatmaps were made using Deeptools ComputeMatrix (options: reference-point; missingDataAsZero) and PlotHeatmap.

### RNA-seq data processing

Reads were mapped using HISAT2 to the dm6 genome assembly. Mapped reads were then filtered for map quality (SAMtools(*65*)).

Differential analysis was performed using DESeq2 with cutoff: padj < 0.01.

For comparison of ATAC-seq and RNA-seq, all ATAC-seq peaks were assigned to the nearest gene that is expressed in either wing or haltere imaginal disc (count > 50). For each gene the single ATAC peak with the lowest p-value determined by DESeq2 differential analysis, and the (W/H) -Log_10_P as determined by DESeq2 (described above) was compared between peaks associated with differentially expressed vs non-differentially expressed genes.

### ChIP-seq data processing

Reads were mapped using Bowtie2 to the dm6 genome assembly. Mapped reads were then filtered for map quality (SAMtools (*65*)) and duplicates (Picard MarkDuplicates). Peaks were called using MACS2 (*67*). Genome-track files were created using Deeptools (BamCoverage; RPKM normalization).

For comparison of Sd binding in wing and haltere, differential analysis was performed using DiffBind (*68*) (FDR < 0.05 for significance cutoff).

### Motif analysis

*De novo* motifs were discovered using Homer (*69*) (findmotifsgenome.pl). For ATAC-seq data the entire peak was used to search for enriched motifs (option: -size given). For ChIP-seq a default 200bp window around the peak center was used. To center peaks around the best match to the de-novo motif (Fig. 2E, F) the annotatepeaks command was used (option: -mbed) to generate the location of the motif.

---

**Fig. S1.**
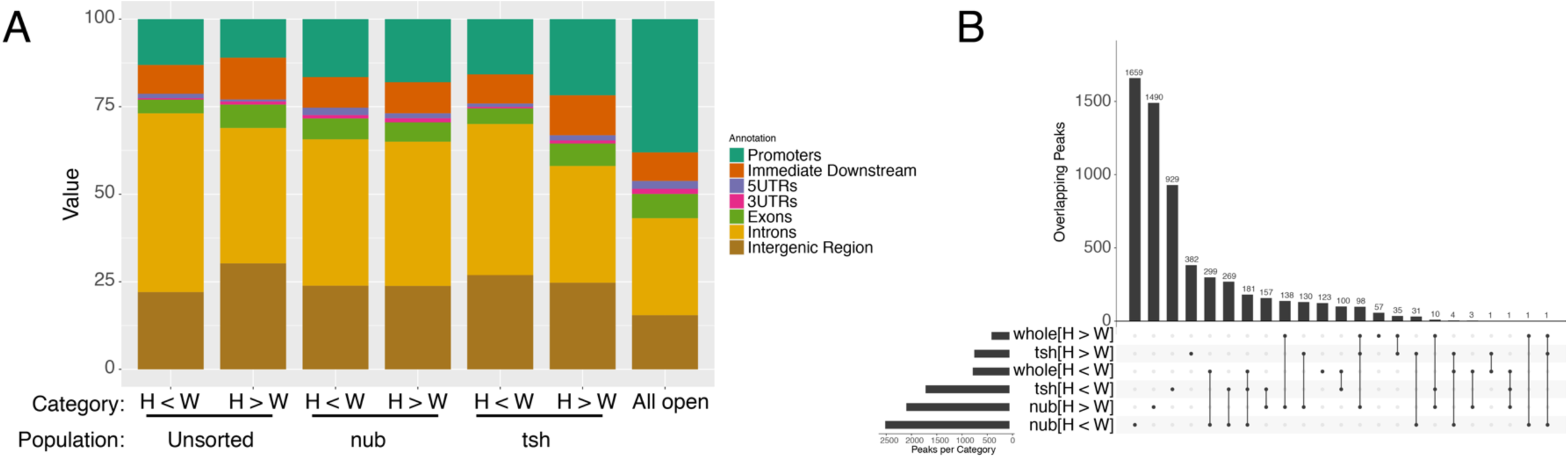
Annotation of ATAC-seq peak sets. A. Histogram showing the distribution of ATAC-seq peaks relative to various genomic regions. B. UpsetR plot showing overlap between differential ATAC-seq peaks.

**Fig. S2.**
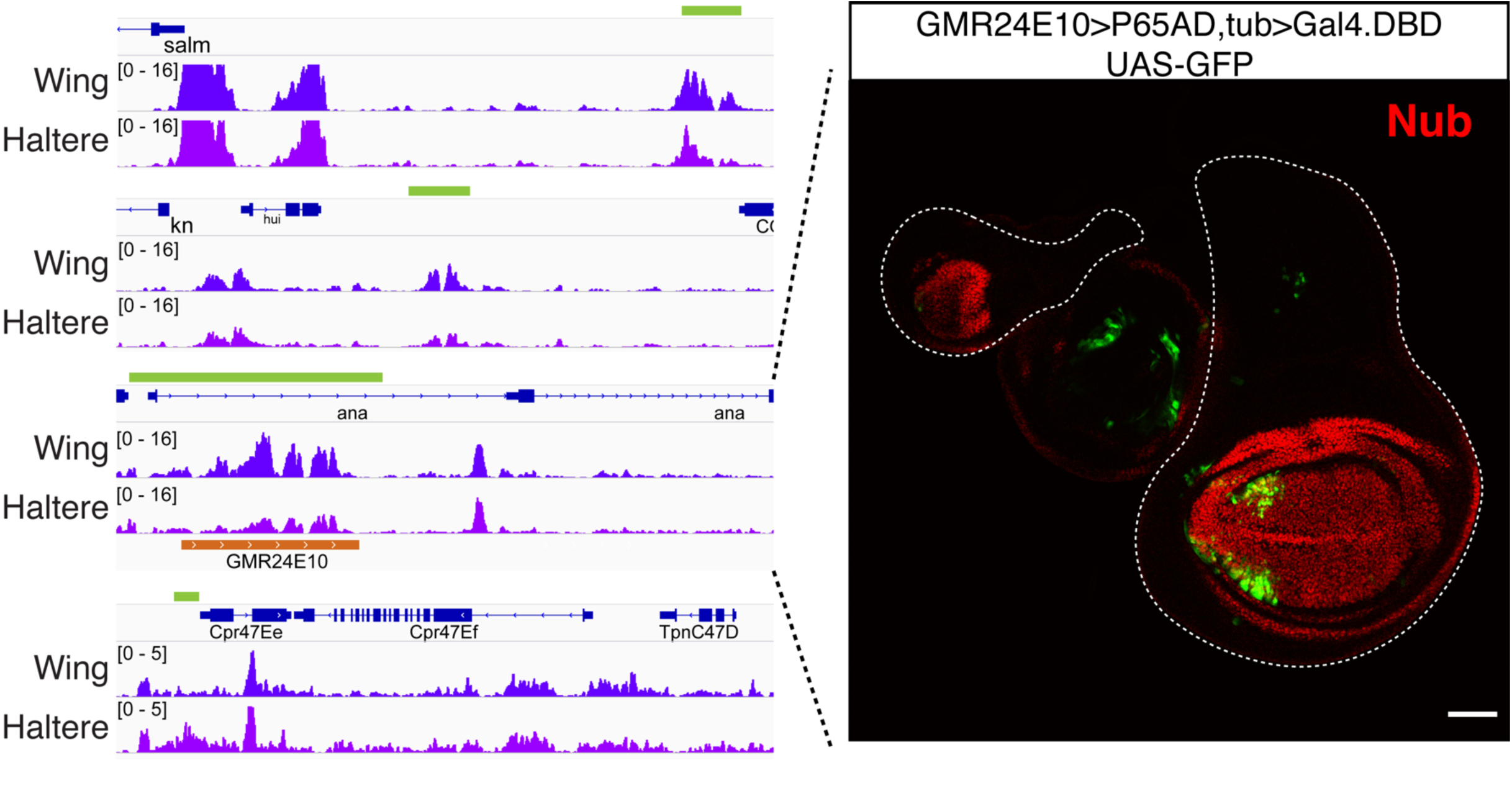
Known Ubx target CRMs are differentially accessible in wing and haltere. A. Genomic tracks of unsorted ATAC-seq wing and haltere for previously identified Ubx targets in the haltere. Green bars represent the originally defined enhancer boundaries. B. Reporter expression of *ana-spot* CRM using a enhancer fragment generated by (*70*) that recapitulates previously described pattern(*17*). See Fig.1G-H for expression of *sal1.1* and *KnW* fragments. For expression of the Cpr47ee enhancer see reference(*17*).

**Fig. S3.**
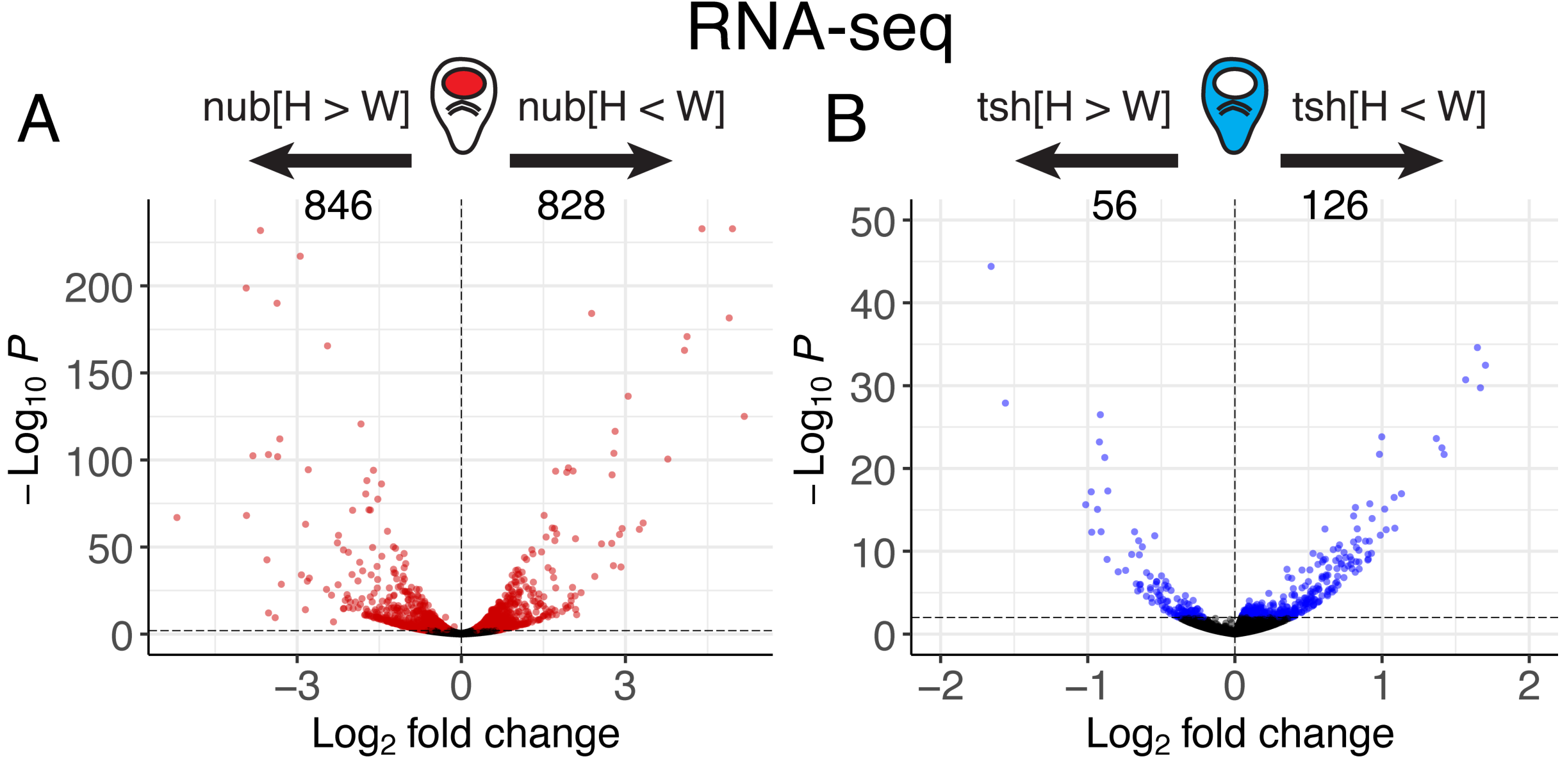
Differential RNA-seq analysis. A-B. Volcano plots displaying results of differential RNA-seq analysis between wing and haltere discs for *nub+* (A) and *tsh+* (B) populations. Colored points are based on a significance threshold of padj<0.01 (DESeq2).

**Fig. S4.**
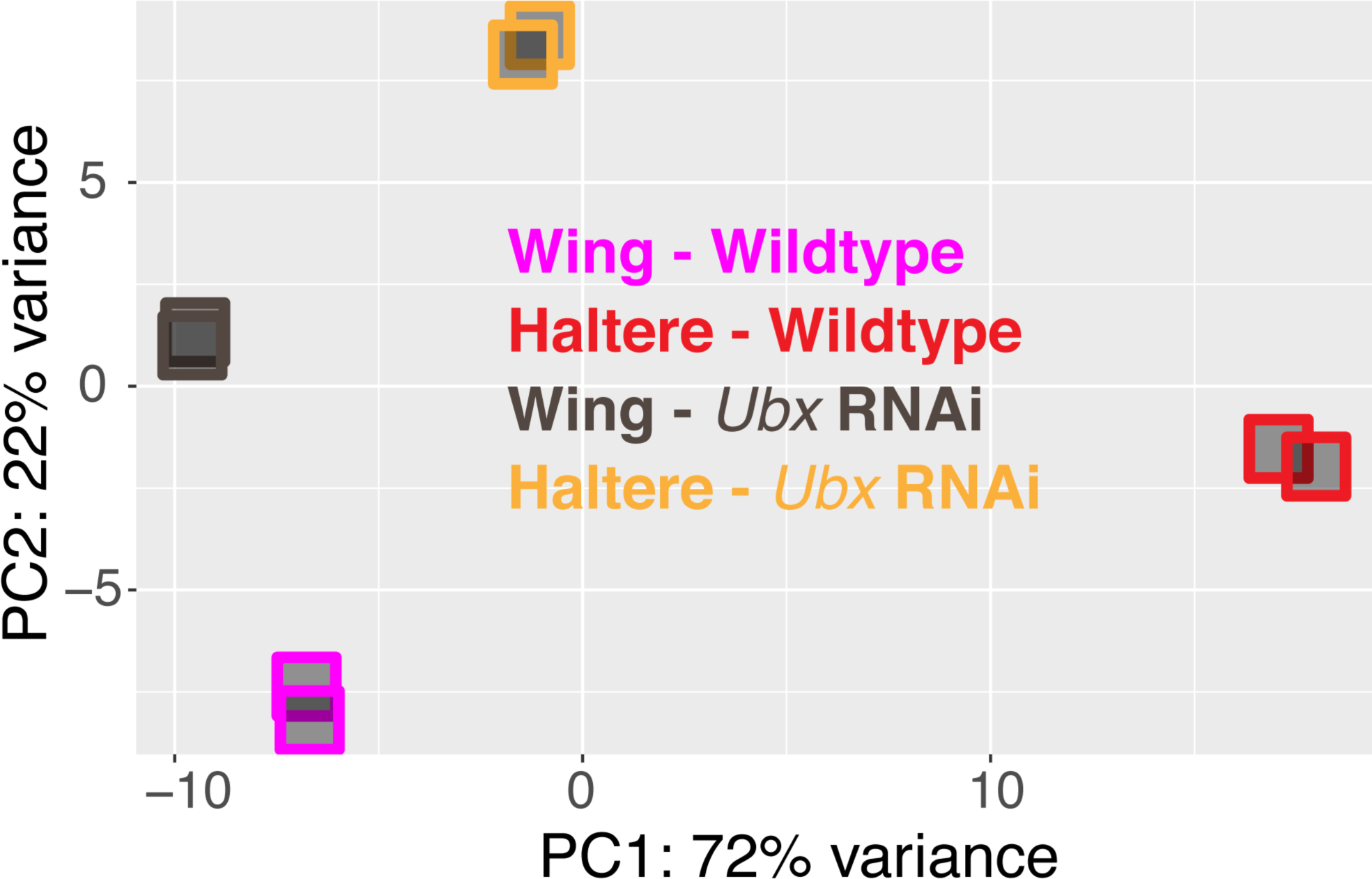
PCA of wild-type and *Ubx* knockdown ATAC datasets. Principal component analysis (PCA) for sorted *nub+* ATAC: wing and haltere wild-type (purple and red, respectively) and expressing Ubx-RNAi (black and orange, respectively).

**Fig. S5.**
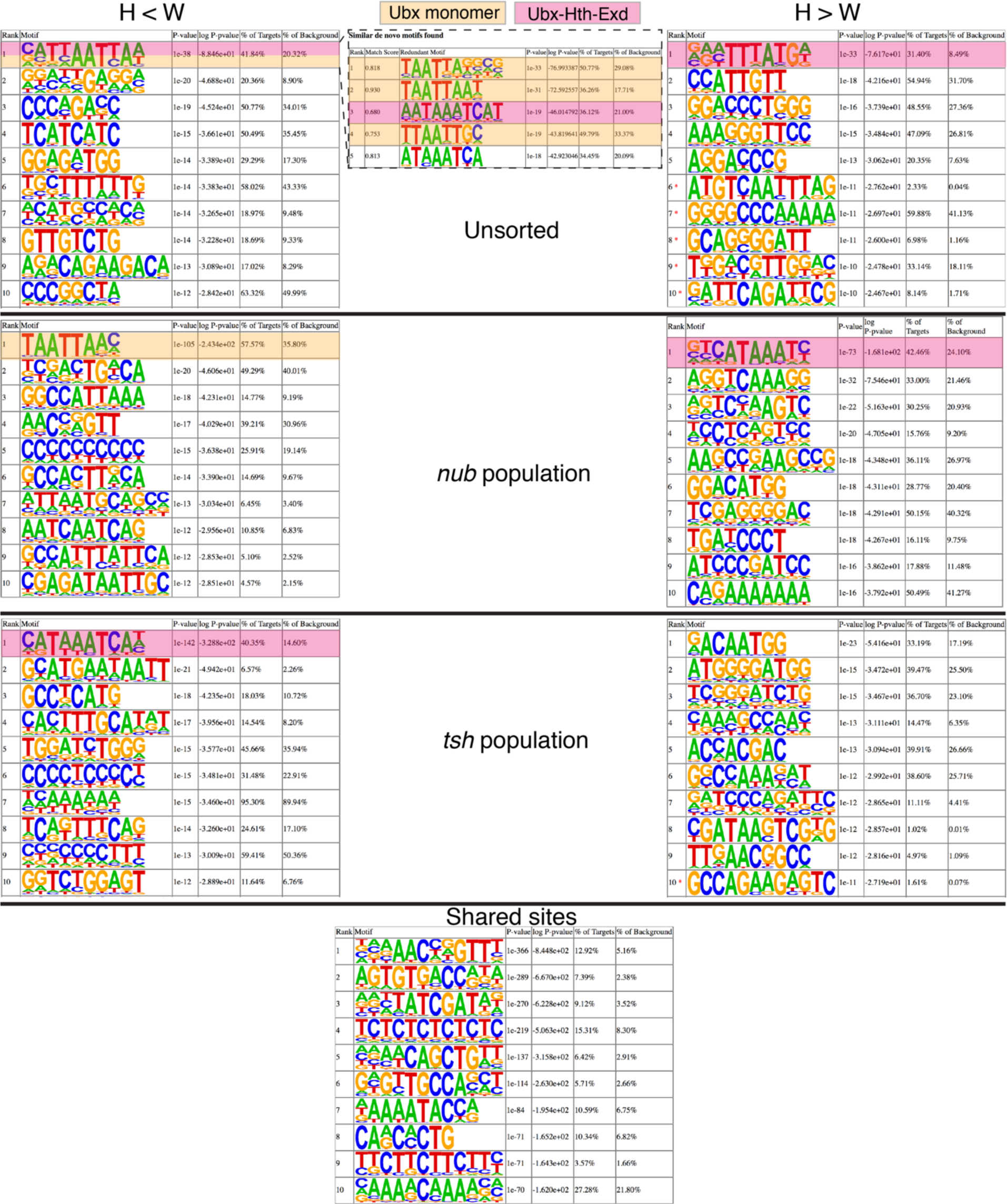
ATAC-seq motif enrichment. Top 10 *de novo* motifs identified from ATAC-seq peaks from differential unsorted (additional panel in H<W category shows similar motifs grouped together for the 1^st^ ranked position), differential sorted *nub+*, differential sorted *tsh+*, and shared ATAC-peaks. Boxes on the left and right represent regions that decrease and increase accessibility in the haltere datasets, respectively. Red asterisks represent motifs that do not reach statistical significance (p-value < 1e-12). Ubx monomer and Ubx-Hth-Exd complex motifs are labeled in orange and red, respectively.

**Fig. S6.**
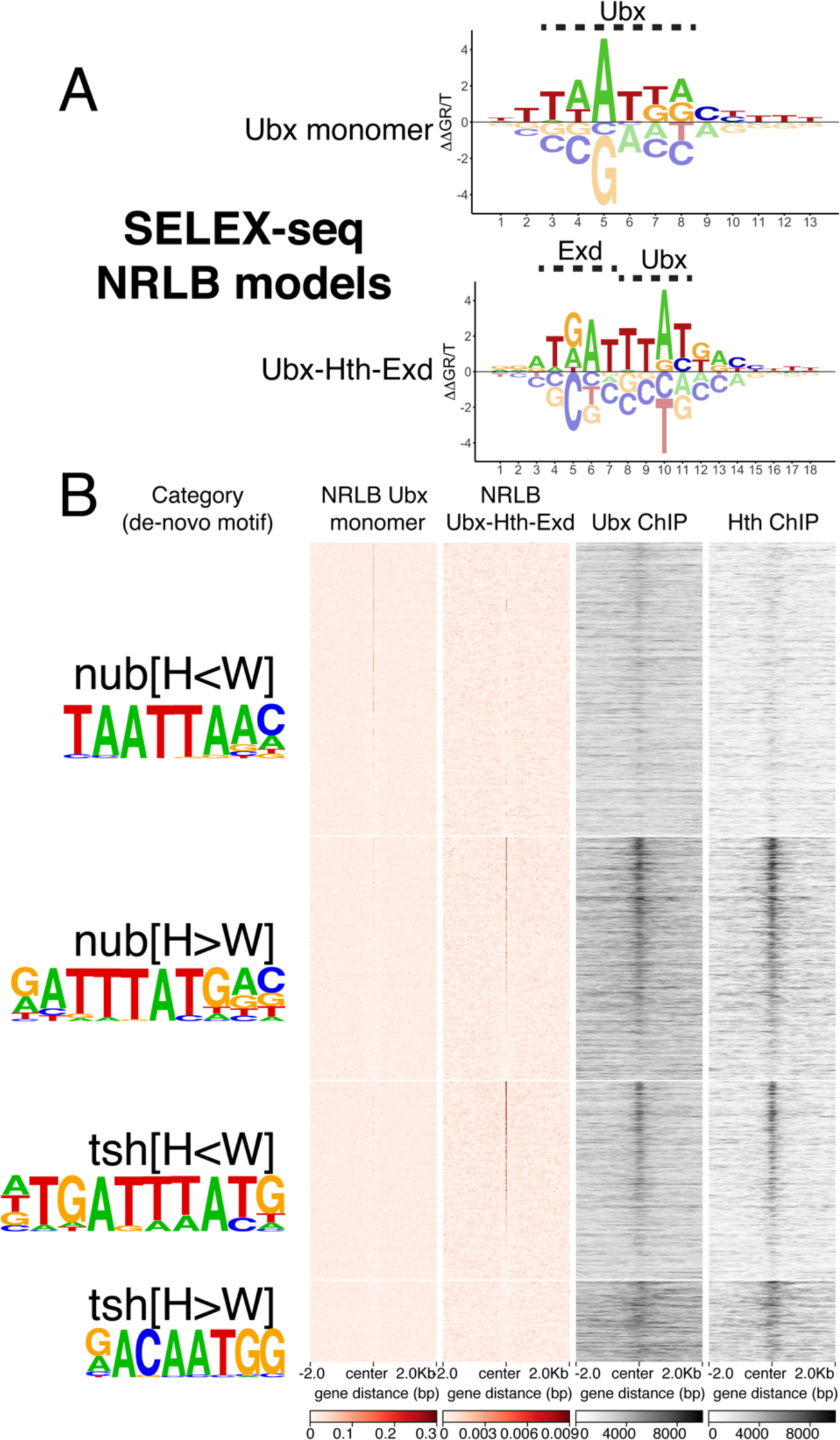
NRLB-based scoring of the four ATAC-seq categories. A. Energy logos for Ubx monomer (Ubx isoform IVa) and Ubx-Hth-Exd (with Ubx isoform IVa) derived from NRLB modeling of SELEX-seq data ((*34*), https://github.com/BussemakerLab/NRLB). Nucleotides that interact with Ubx or Exd are labeled above. B. Heatmaps sorted as in Fig. 2E (centered on the *de-novo* motif shown to the left) and scored with NRLB models (red columns) confirm that three of the four ATAC-seq categories contain binding sites for Ubx or Ubx-Hth-Exd, as predicted by the *de novo* motif analysis. Associated whole haltere ChIP-seq heatmaps for Ubx and Hth are also shown (as in Fig. 2E).

**Fig. S7.**
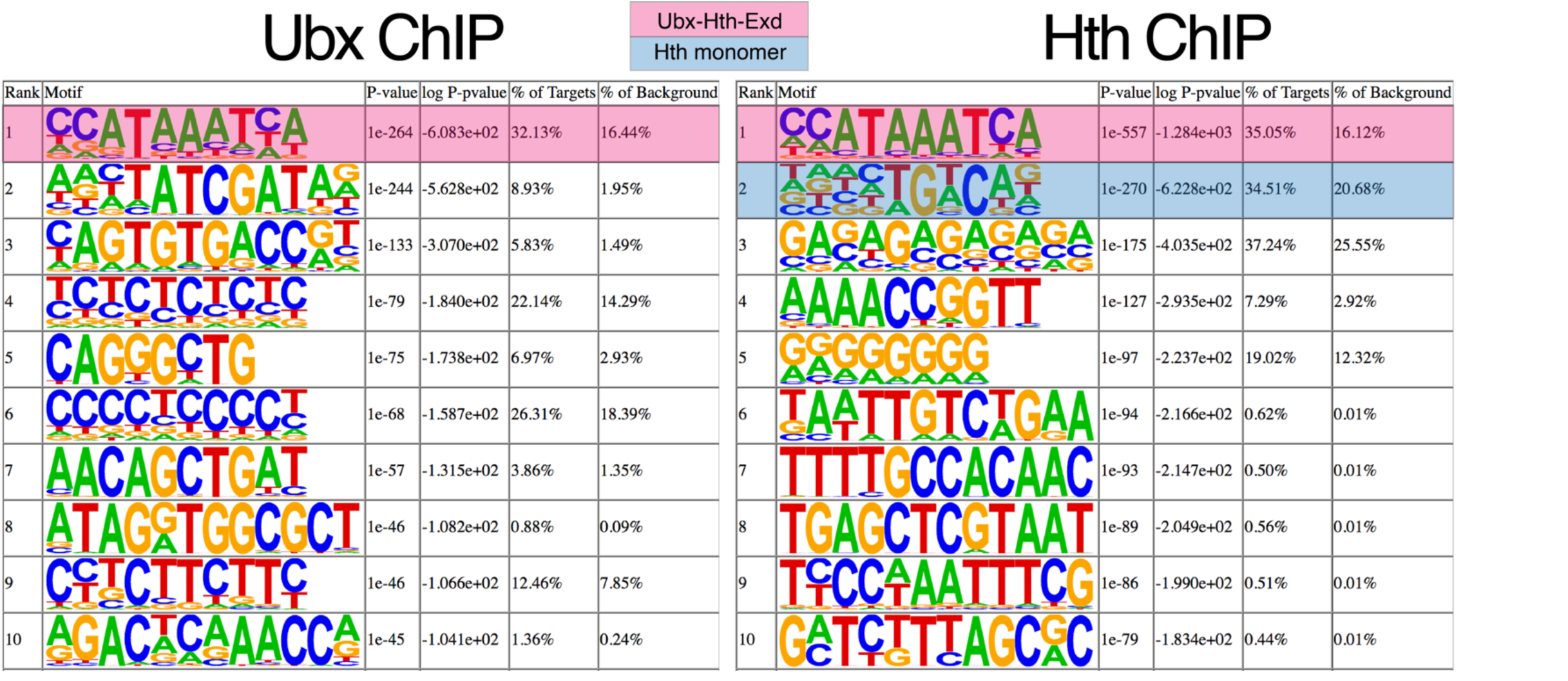
Ubx and Hth ChIP-seq motif enrichment. *de novo* motifs identified from ChIP-seq peaks of Ubx (Left) and Hth(Right) identified in haltere imaginal discs. Ubx- Hth-Exd and Hth monomer motifs are labeled in red and blue, respectively.

**Fig. S8.**
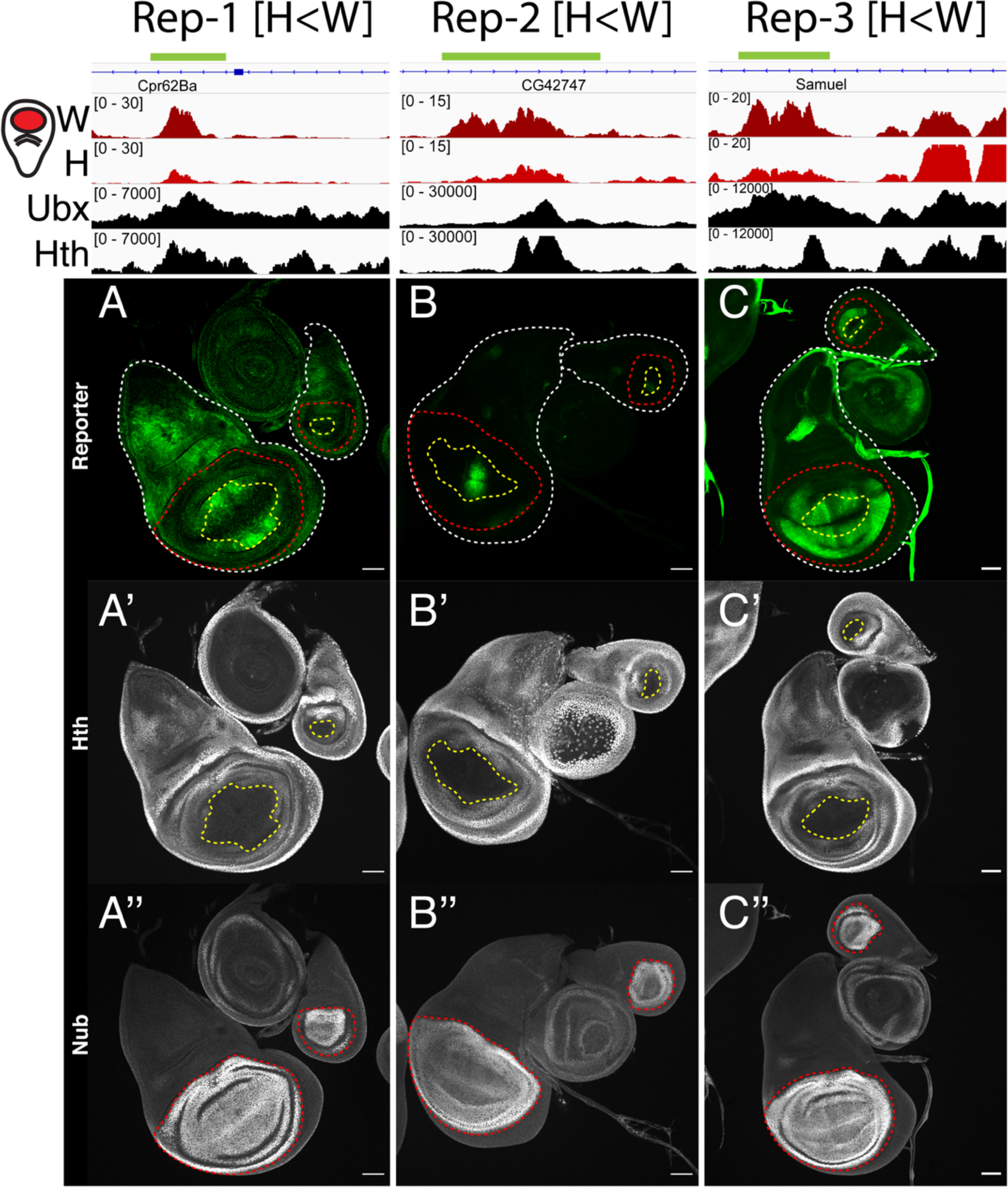
nub[H<W] reporters. A-C. Reporter expression of CRMs from the nub[H<W] category. For this and subsequent reporter gene figures, the top two genome browser tracks show the ATAC-seq signal in the *nub+* wing (W) and haltere (H) imaginal discs.

**Fig. S9.**
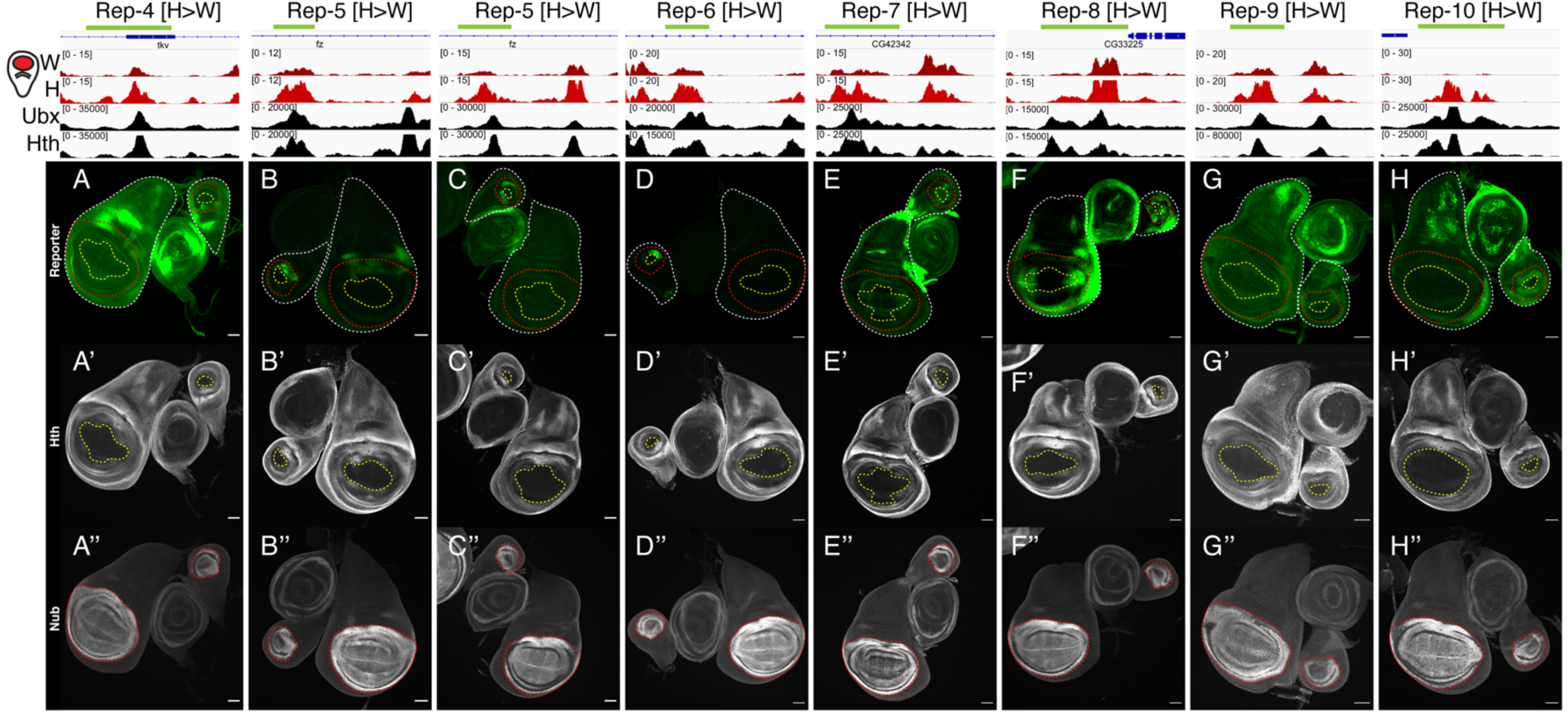
nub[H>W] haltere-only reporters. A-H. Reporter expression of CRMs from the nub[H>W] haltere-specific category.

**Fig. S10.**
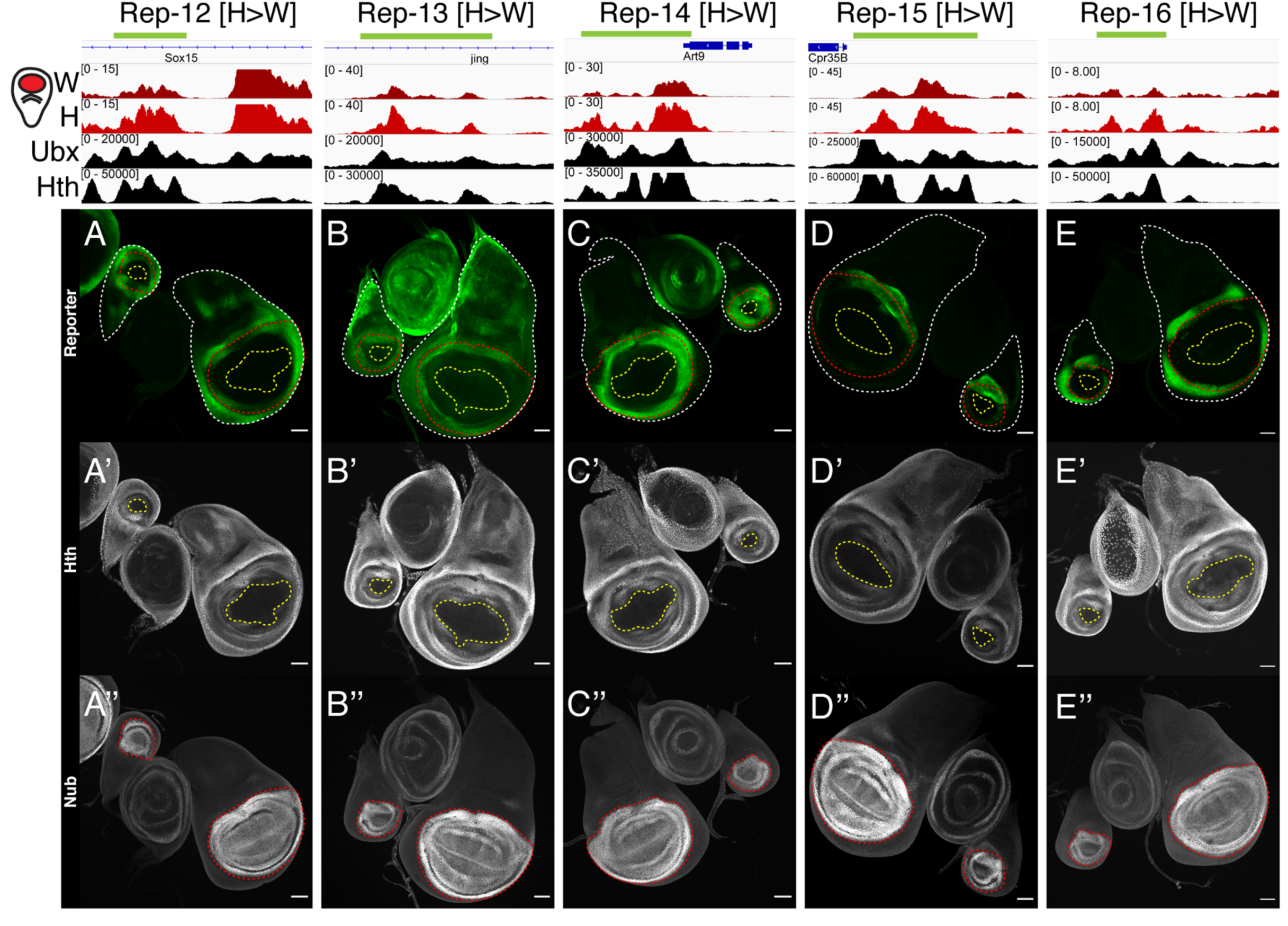
nub[H>W] haltere-expanded reporters. A-F. Reporter expression of CRMs from the nub[H>W] haltere-expanded category.

**Fig. S11.**
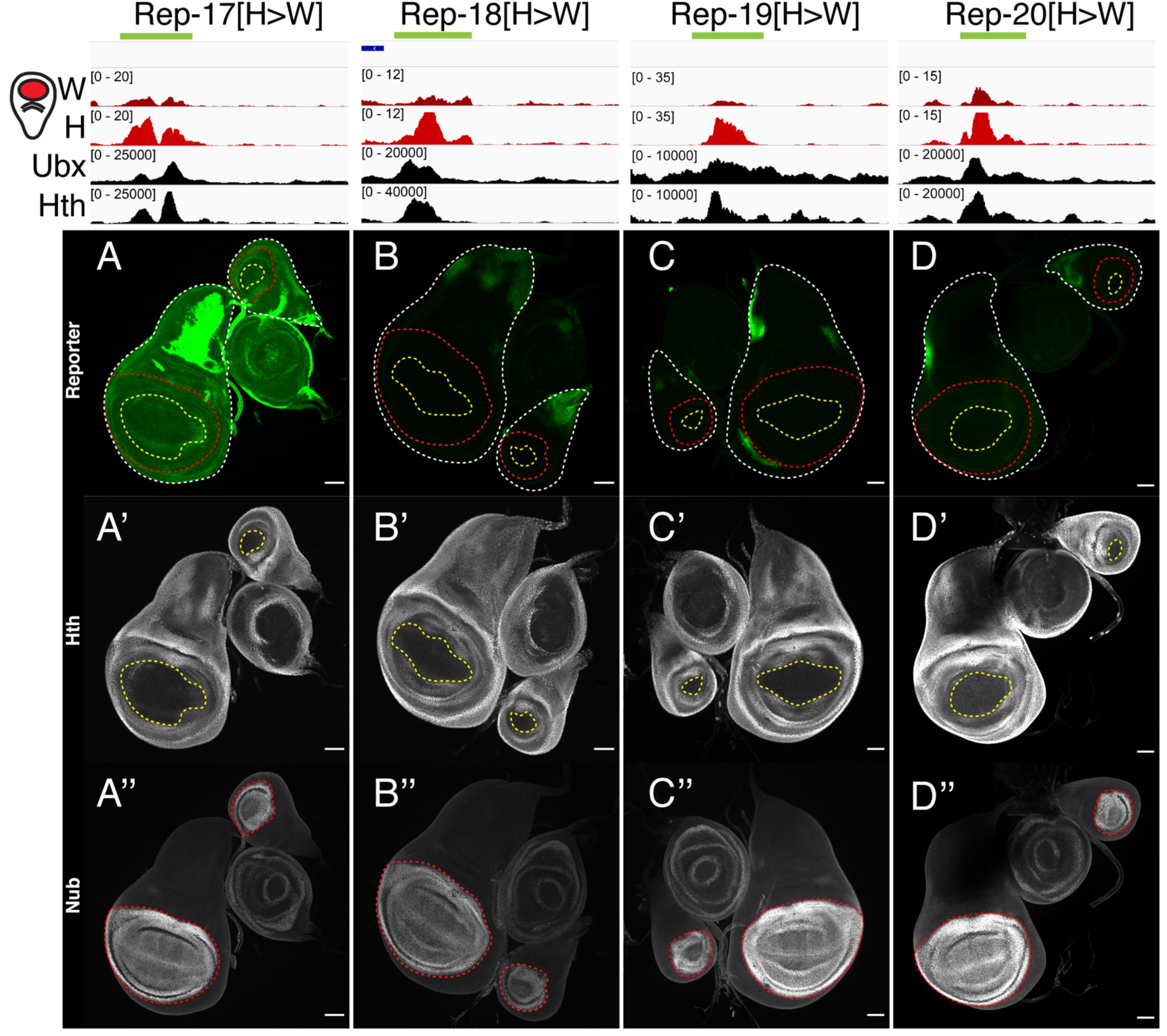
nub[H>W] No-nub activity reporters. A-D. Reporter expression of CRMs from the nub[H>W] No-nub activity category.

**Fig. S12.**
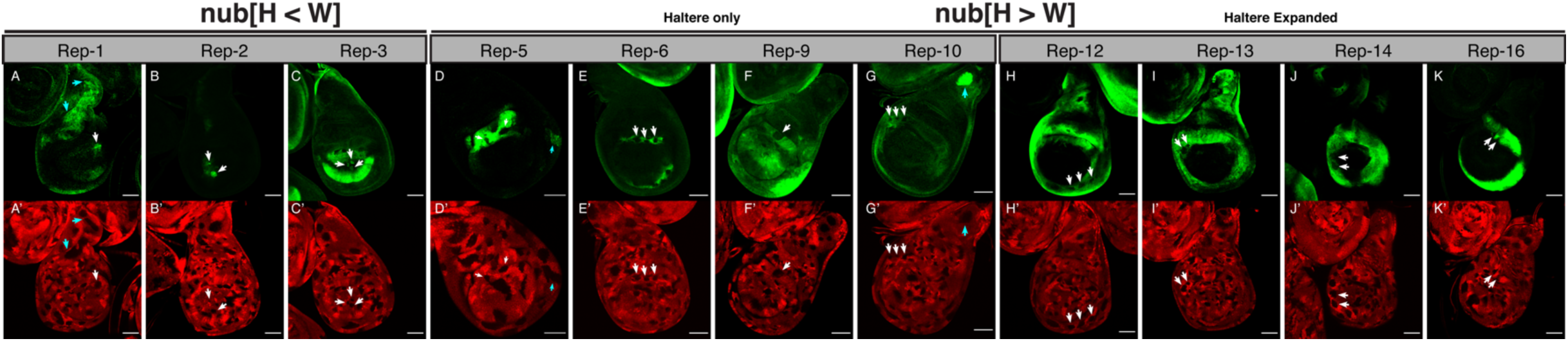
Reporter gene activity in *Ubx-* mitotic clones. A-K. *Ubx*-null clonal analysis of reporter expression. Clones are marked by the absence of RFP (A’-K’). Clones in *nub+* region are marked by white arrows, and clones derepressed in *tsh+* region are marked by cyan arrows. Note that only a subset of clones reveal a change in expression, consistent with the existence of additional regulatory inputs into these CRMs. For comparison, the wing disc expression patterns are shown in Fig.s S8-S10.

**Fig. S13.**
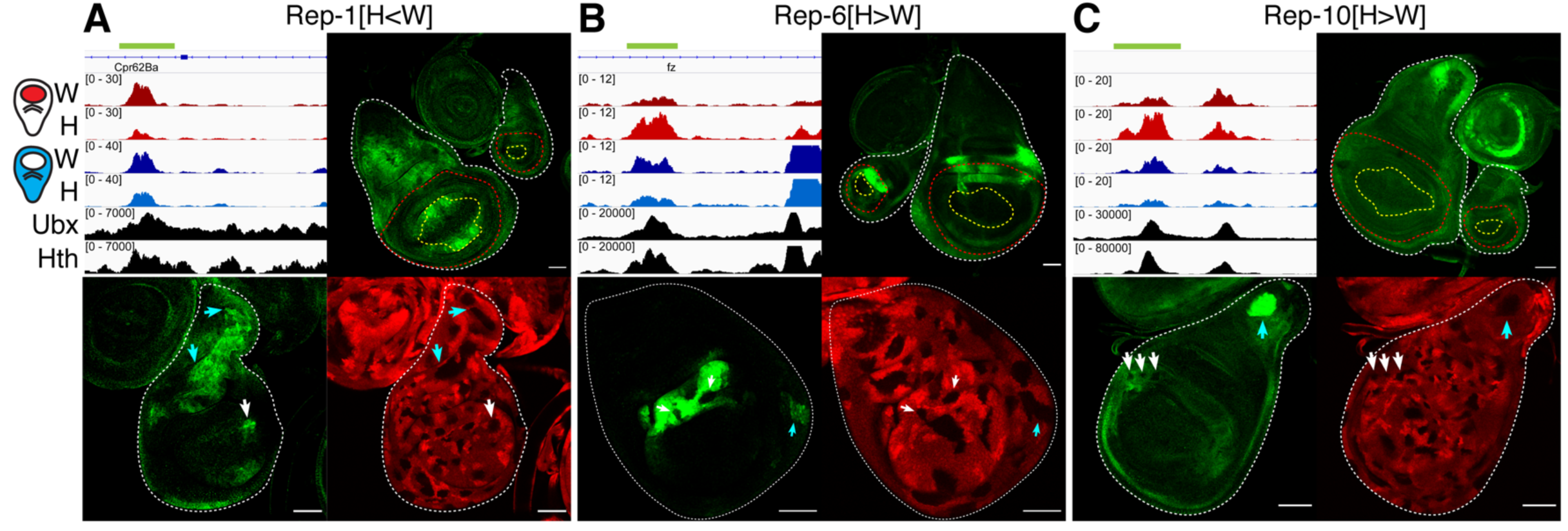
Reporters responsive to Ubx in both the *nub+* and *tsh+* domains. A-C. Reporters that are repressed by Ubx in the *tsh+* population in addition to regulation in the *nub+* cells (green). The upper left panels show genomic tracks for *nub+* ATAC-seq wing, *nub+* ATAC-seq haltere, *tsh+* ATAC-seq wing, *tsh+* ATAC-seq haltere, Ubx ChIP, and Hth ChIP; the upper right panels show wing and haltere disc expression patterns for the reporter genes, and the bottom panels show *Ubx* null somatic clones in the haltere marked by the absence of RFP (bottom-right panel). A subset of clones are marked by arrows, and the position indicated by white (*nub+*) or Cyan (*tsh+*).

**Fig. S14.**
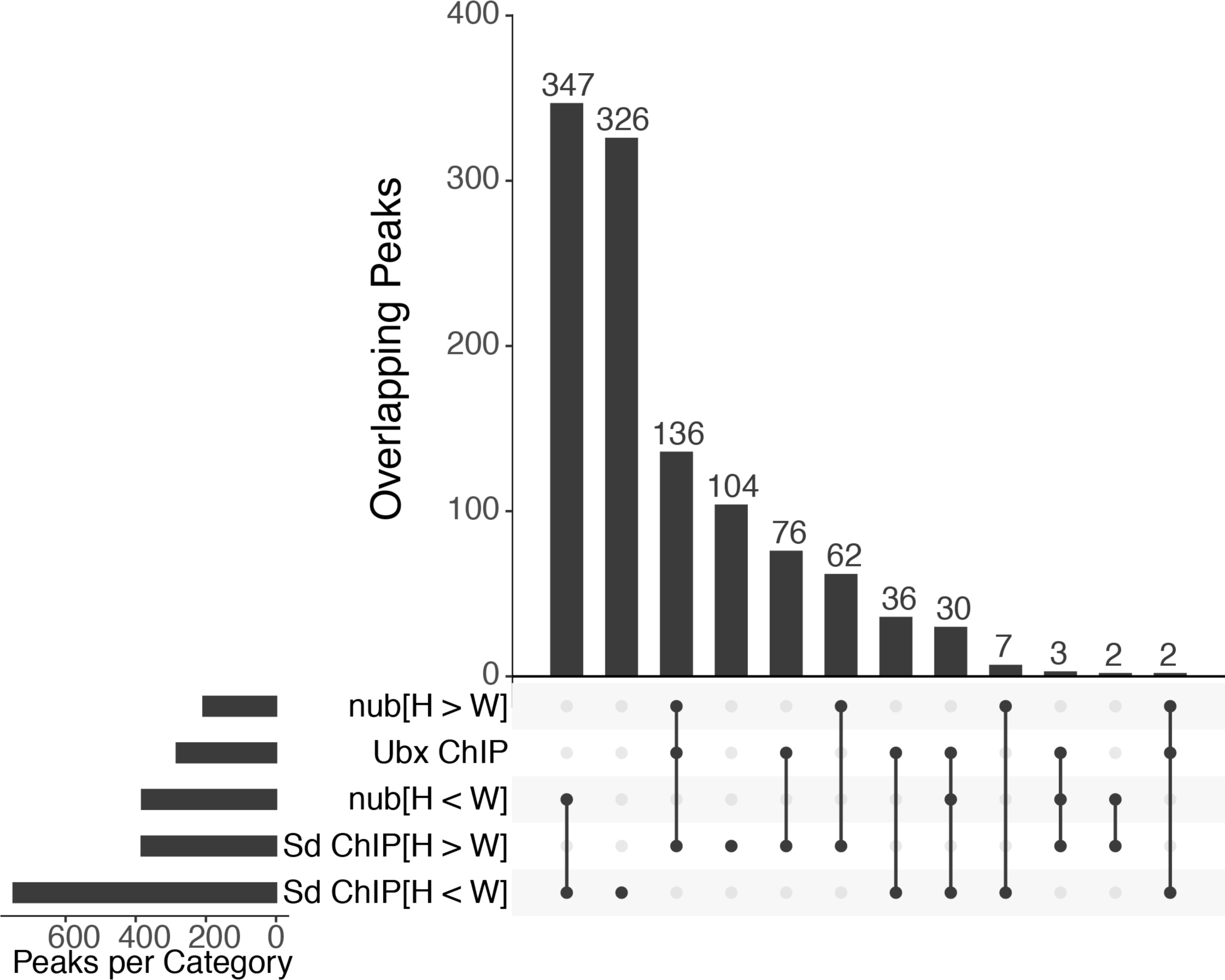
Overlap of tissue-specific Sd binding. UpsetR plot showing overlap between the Ubx ChIP-seq, differential ATAC-seq (nub[H<W] or nub[H>W]), and differential Sd ChIP-seq peak sets (sd[H < W] or sd[H > W]).

**Fig. S15.**
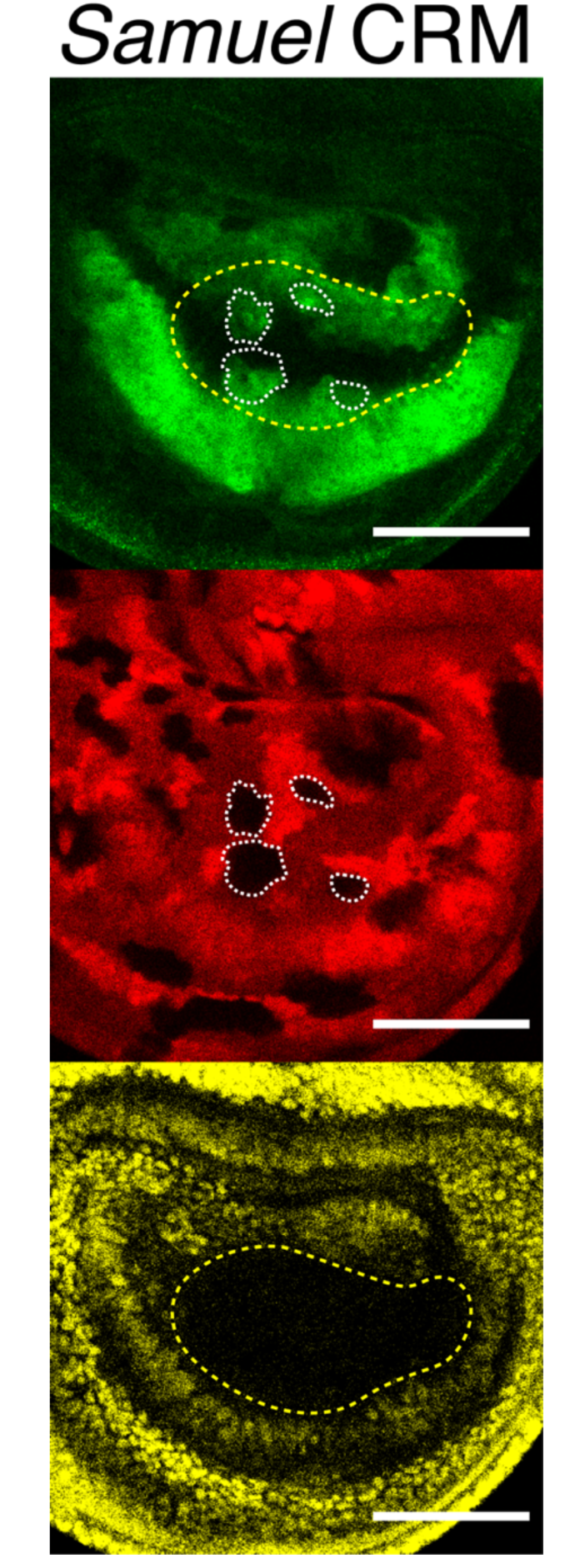
Samuel CRM expression in *Ubx-* clones. Additional example of *Ubx* null clones and *Samuel* CRM (Rep-3 nub[H<W]) activity. Only a subset of clones within the pouch show de-repression, while regions where wild-type activity of the CRM in the is similar to wing (dorsal and ventral-most positions) are unaffected.

**Table S1.**
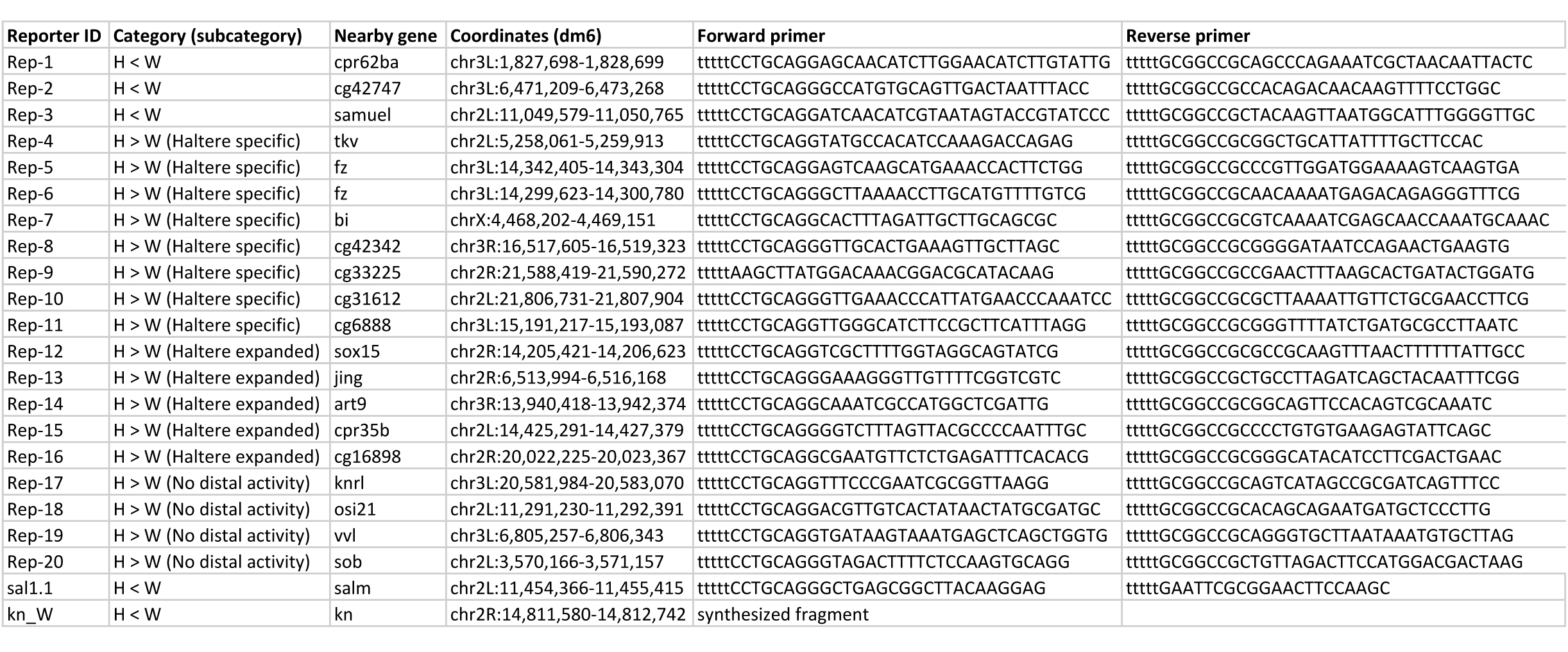
Transgenic reporters

